# Resident liver macrophages buffer dietary and circulating cholesterol to maintain hepatocyte homeostasis

**DOI:** 10.64898/2026.11.13.766502

**Authors:** Matthew Hamilton, Syed Monzur Morshed, Madison Chlebowski, Celia E. Shiau

## Abstract

Cholesterol metabolism is viewed largely through the lens of hepatocytes, leaving the roles of other liver cells unclear. Using high-resolution live imaging and cholesterol tracing in intact zebrafish, complemented by genetic and clodronate-mediated macrophage depletion and single-cell analysis, we identify liver-resident macrophages (LMs) as cellular buffers of circulating and dietary cholesterol. LMs directly take up cholesterol, and their depletion redirects cholesterol to hepatocytes, which suppress cholesterol synthesis, increase bile-acid synthesis and efflux, and activate metabolic-stress and immune programs, including hepatokine *lect2* induction. These responses are heightened by elevated dietary cholesterol and occur without overt inflammation or altered liver architecture. LMs partition cholesterol through *abca1b*-dependent export and *cetp*-mediated lipoprotein remodeling. Loss of *abca1b* causes cholesterol sequestration in LMs and reduced cholesterol-processing demand on hepatocytes, while LM depletion eliminates macrophage-derived *cetp* and reduces hepatocyte *cetp*. Together, these findings reveal the innate immune system as integral to normal metabolic physiology, with LMs buffering cholesterol to maintain hepatocyte homeostasis.

## Introduction

The liver is a central regulator of whole-body homeostasis, integrating metabolism, detoxification, nutrient distribution, and synthesis of essential circulating proteins while coordinating systemic glucose and lipid balance^1–3^. At the same time, it functions as a major immunological interface between the intestine and the circulation, continuously exposed to dietary metabolites, microbial products, and blood-born signals arriving through the portal vein. The liver serves as both a metabolic hub and an immune sentinel, requiring tight coordination between parenchymal and immune cell populations to preserve local and systemic homeostasis^1,2^. Nutrients and microbial-derived factors absorbed from the intestine enter the liver directly through the portal circulation, enabling hepatocytes to process dietary lipids including cholesterol while simultaneously filtering potential pathogens and microbial products from the gut^1,2^. In parallel, systemic circulation from peripheral tissues is continuously routed through the hepatic artery to the liver, allowing hepatocytes to integrate metabolic feedback from the rest of the body and adjust endocrine, biosynthetic, and catabolic outputs accordingly^1–3^. Thus, the liver operates as a bidirectional gateway linking intestinal nutrient flux with systemic metabolic regulation.

Among hepatic immune populations, resident macrophages constitute one of the largest tissue macrophage compartments in the body^4–6^. Although classically viewed as sinusoid-associated Kupffer cells specialized for immune surveillance and pathogen clearance, recent studies have revealed substantially greater heterogeneity in liver-resident macrophages (referred herein as liver macrophage, LM), expanding on their identity, localization, and molecular state in the mammalian liver^4–6^. Although hepatocytes are recognized as the central cellular hub for hepatic nutrient metabolism, how nutrient flux is partitioned among distinct liver cell populations, and whether resident macrophages actively contribute to this process under physiological conditions remains poorly understood.

To address this gap, we leveraged the zebrafish as an ideal vertebrate model for studying complex physiological questions across cellular, tissue, and organismal scales owing to its accessibility for in vivo imaging, genetic manipulation, molecular interrogation, and physiological analysis, while retaining major conserved metabolic pathways with humans^7–10^. Zebrafish has been widely used to study liver development, regeneration, and lipid metabolism, including fatty acid and lipoprotein biology^7–15^. However, studies of the zebrafish liver have historically focused on embryonic or early larval stages before sustained feeding, or on larvae maintained under confined conditions in petri dishes that impair hepatic maturation. These limitations contributed to a longstanding misconception that the zebrafish liver lacks resident macrophages^7,8,16^ until this study and more recent publications^17,18^.

Here we show that, as in mammals, zebrafish maintain large numbers of resident macrophages in the liver throughout life. Using *irf8* mutants as a model for genetic ablation of macrophages^19–22^, we found disrupted hepatocyte homeostasis particularly after increased dietary cholesterol. Loss of LMs redirects cholesterol toward hepatocytes, increasing its disposal through efflux and bile acid production to maintain cholesterol homeostasis. Simultaneously, hepatocytes in the absence of LMs activate immune and stress responses^23–26^, including the induction of hepatokine *lect2.2*, cellular defense gene *hsp70.3*, viral response gene *isg15*, c-reactive protein gene *crp5*, and complement-associated genes *c4b* and *cfhl5*. Of particular interest, the transcriptional increase of the hormone-like protein LECT2 has been associated with adaptive responses to metabolic stress or immune activation in liver diseases, as well as in stimulating cellular growth and proliferation of bone and cartilage cells^23,24,27^. In *irf8* mutants, we found a small number of residual immature myeloid cells to be strikingly distinct from functional liver macrophages and unable to properly take up and process cholesterol from either circulation or diet, thereby effectively making these livers LM-depleted. Strikingly, these cells ectopically induced both paralogs of the hepatocyte hormone-like LECT2, *lect2.2* and *lect2.1*, irrespective of diet, potentially amplifying hepatocyte stress hormone signaling. Furthermore, the increased cholesterol in hepatocytes does not appear to cause lipid droplet accumulation but may instead increase the pool of free cholesterol that promotes bile acid synthesis and efflux in *irf8* mutants. These findings suggest that liver macrophages continuously buffer physiological cholesterol levels under normal conditions. Mechanistically, liver macrophages actively shape hepatic cholesterol flux by directly capturing and processing cholesterol, exporting cholesterol through macrophage-restricted *abca1b*, and contributing to *cetp*-associated lipoprotein remodeling that regulates the hepatic influx and partitioning of cholesterol from dietary and peripheral sources. Together, our findings establish resident liver macrophages as central regulators of metabolic and immune homeostasis in the healthy liver.

## Results

### Mapping and characterizing resident macrophages reveal their biliary localization in the zebrafish liver

We established growth and feeding conditions using one-liter tanks to develop fully mature zebrafish larvae through two weeks post-fertilization in the laboratory (Supplementary Fig. 1, Supplementary Movie 1). This approach was critical to overcome deficient liver maturation from larval growth confined to 10 cm petri dishes with little to no feeding past 5 days post-fertilization (dpf) that has been common practice in the field. Appropriately grown larval zebrafish revealed colonization of resident macrophages within the liver of larvae beginning at 6–7 dpf which have grown to a minimum standard length of 4 mm (Fig. 1 and Supplementary Fig. 2). By 8 dpf, a substantial number of macrophages have taken residence in the liver mostly along the biliary network with some intimately associated with the vasculature (Fig. 1b). As zebrafish reaches 10 to 12 dpf, there are on average more than 20 liver macrophages per liver and this number steadily increases through 17 dpf and into adulthood (Fig. 1b-e and Supplementary Fig. 3).

**Fig. 1.**
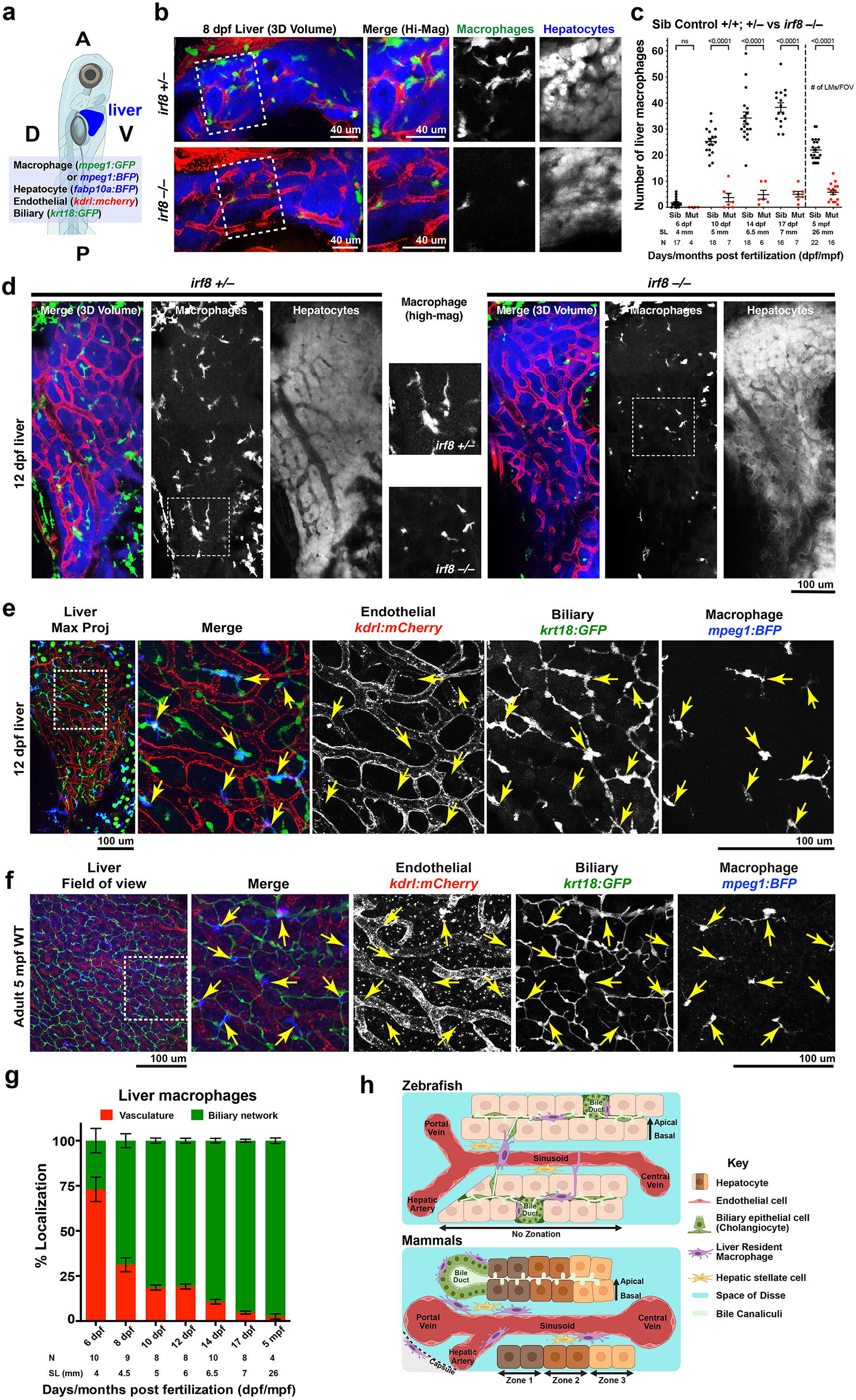
Liver-resident macrophages (LMs) persist along the biliary network throughout zebrafish life and are depleted in *irf8* mutants. **a** Schematic showing the orientation of larval zebrafish for in vivo confocal imaging of the liver (blue). Transgenic reporters for macrophages, hepatocytes, endothelial cells, and biliary epithelial cells were used in different combinations to map LM localization. A, anterior; P, posterior; D, dorsal; V, ventral. **b** Representative confocal images at 8 dpf showing the emergence and morphology of LMs (GFP+) within or adjacent to the vasculature (red); the first LMs are detectable at 6 dpf. Right, higher-magnification views of dotted boxed regions showing merged and individual macrophage and hepatocyte channels. In *irf8* mutants, typical elongated LMs are absent and replaced by few small, rounded residual *mpeg1*+ cells. **c** Quantification of LM abundance from 6–17 dpf and in adults, showing progressive detection of LMs throughout development and adulthood, whereas residual small-sized *mpeg1*+ cells remain rare in *irf8* mutants. Data are aligned with fish size based on standard length (SL); N, independent animals analyzed. **d** Representative confocal images at 12 dpf showing redistribution of LMs from the vasculature into the liver parenchyma. Higher-magnification views highlight the distinct morphology of control LMs, which are elongated with elaborate processes, compared with the small, rounded residual *mpeg1*+ cells in *irf8* mutants. **e** Representative confocal images at 12 dpf showing the spatial relationship of LMs with the vascular and biliary networks. LMs (yellow arrows) show clear cellular overlap with biliary epithelial cells, but no alignment with blood vessels. Individual channels and merged images are shown. **f** Representative confocal images of adult zebrafish liver showing the same association of LMs (yellow arrows) with the biliary network (green), rather than the vasculature (red), is maintained into adulthood. **g** Quantification of LM localization relative to the vascular and biliary networks, showing an early vascular association that shifts toward predominantly biliary by 8-10 dpf and persists throughout development into adulthood. **h** Schematic comparing the spatial and cellular organization of zebrafish and mammalian liver architecture. Statistical significance was determined using a two- tailed Students *t*-test. Data are presented as mean ± SEM.

Consistent with emerging mammalian single-cell and histological studies demonstrating macrophage populations beyond sinusoidal Kupffer cells, including macrophages associated with bile ducts, the capsule, and the central vein^4^, we found zebrafish liver macrophages to localize predominantly along the biliary epithelial network while extending highly stable processes into and around hepatic sinusoids (Fig. 1 and Supplementary Figs. 3,4, Movies 2,3). Over their developmental time course, liver macrophages increasingly shift from vascular to biliary localization in the zebrafish liver (Fig. 1f). While mammalian liver is organized in lobules with a clear portal triad and central vein structure whereby the bile ducts and vasculature form organized regular patterns that are non-overlapping, zebrafish liver lacks such lobular structure and instead has two separate vasculature and biliary cellular networks that are interweaved (Supplementary Fig. 5)^15,28,29^. The sinusoidal blood vessels occupy larger volumes interspersed between hepatocytes on the basal side while the biliary epithelial cells, identified using either keratin-18 (*krt18*) reporter^30^ or Anxa4 (2F11) antibody^28,29^ labeling, form largely preductule structures that take up significantly thinner spaces between hepatocytes on the apical side (Supplementary Fig. 5 and Fig. 2). Despite differences in the liver tissue organization, the biliary localization of zebrafish LMs closely resembles mammalian hepatic lipid-associated macrophages positioned near the bile duct^4,5^, whose functions remain elusive (Fig. 1g).

**Fig. 2.**
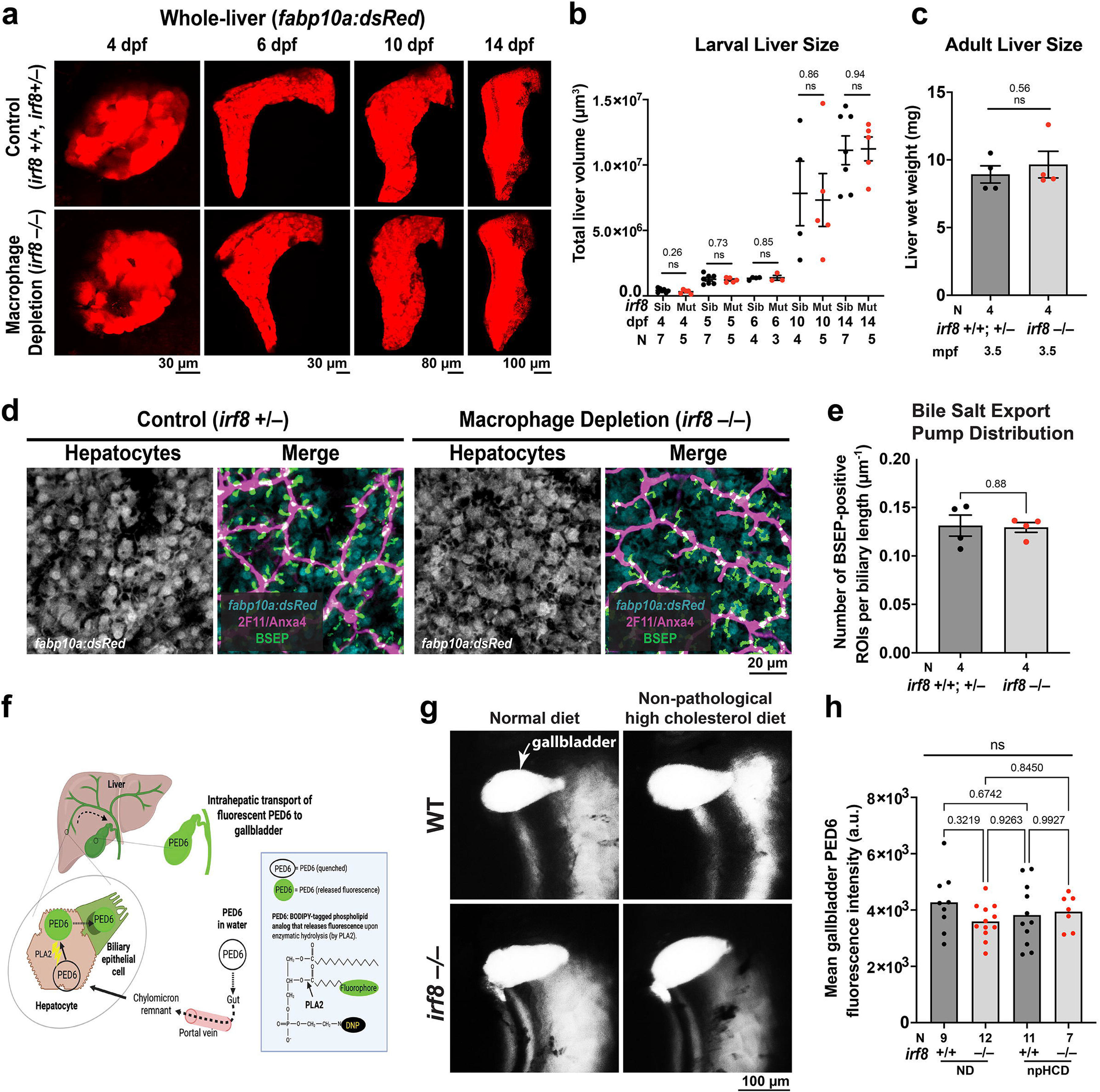
Liver macrophage depletion does not alter liver growth, biliary architecture, or intrahepatic transport. **a** Representative confocal images of whole livers expressing the hepatocyte reporter *Tg(fabp10a:dsRed)* at indicated developmental stages, showing comparable liver morphology and size between *irf8* mutants and control siblings. Images were segmented in Imaris to isolate liver fluorescence from extraneous signal for tissue-volume quantification. **b** Quantification of Imaris-generated 3D surface reconstructions of whole livers shows comparable liver volumes between control siblings and *irf8* mutants from 4–14 dpf. **c** Wet weight of dissected adult livers showing no difference between *irf8* mutants and control siblings. **d** Representative confocal images of the hepatic biliary network immunolabeled for 2F11 (biliary epithelial cells) and bile salt export pump (BSEP) at 17 dpf. **e** Quantification of BSEP-positive regions of interest (ROIs) normalized to total biliary network length, showing comparable BSEP distribution in *irf8* mutants and control siblings at 17 dpf. **f** Schematic of the PED6 assay used to assess hepatobiliary transport in vivo. **g** Representative fluorescence stereomicroscope images following PED6 exposure, showing comparable accumulation of activated fluorescent PED6 in the gallbladder of control siblings and *irf8* mutants. Statistical significance was assessed using a two-tailed Student’s *t*-test. Data are presented as mean ± SEM; N, number of animals analyzed; sib, *irf8* heterozygous or wild type siblings; mut, *irf8* mutants; dpf, days post fertilization; mpf, months post fertilization; ND, normal diet; npHCD, non-pathological high-cholesterol diet.

Time-lapse imaging in the mature living larval zebrafish without perturbations allows us to study the normal cellular dynamics and behaviors of LMs in detail at homeostasis (Supplementary Fig. 3-4, Supplementary Movies 2-3). LMs can extend extremely fine dynamic processes that protrude into the sinusoidal vessels, often at multiple contact points. Interestingly, their cellular bodies are capable of partially crossing into the blood vessel lumen while they largely remain stationary and embedded in the hepatic parenchyma along the biliary epithelial cells (Supplementary Fig. 3, Supplementary Movie2).

To determine the impact of LM loss on liver physiology, we used the established macrophage-depleted *irf8* mutant model^19^, and conducted a time course analysis of LM counts from their first appearance in the liver through adulthood (Fig. 1b,c). Consistent with previous reports of using *irf8* mutants to analyze brain^19–21^ and intestinal resident macrophages^21,31^, we indeed found a significant depletion of resident macrophages in the liver without effect on other immune cells like neutrophils (Fig. 1 and Supplementary Figure 3). Notably, the few residual *mpeg1*+ cells in *irf8* mutants had markedly weaker *mpeg1* expression and were morphologically smaller and rounder, lacking the extended processes and appearing like immature myeloid cells (Fig. 1b,c).

### Depletion of liver macrophages does not alter hepatic structures but metabolic functions

Considering the abundance of LMs and their alignment with the biliary epithelial cells throughout larval development, we predicted that LMs may contribute to liver growth and structural organization, particularly the biliary network. Contrary to this prediction, a time-course analysis of liver development based on total liver volume from larval stages and total wet weight from adult livers revealed no significant difference in liver size between control siblings and LM-depleted *irf8* mutants across all stages (Fig. 2a–c). Overall liver morphology was also preserved (Fig. 2a). We next assessed biliary architecture by immunohistochemistry using the 2F11 antibody staining for Anxa4, a robust marker of zebrafish biliary epithelial cells^15,29,32^, together with the bile salt export pump BSEP/Abcb11^29,33^ antibody (Fig. 2d,e;Supplementary. Fig. 6). Loss of LMs did not significantly alter biliary network architecture including the biliary canalicular size based on the distribution and volume of BSEP-positive structures, and the biliary network volume, branching, surface area-to-volume, and branch-point spacing, indicating preserved biliary structures (Fig. 2d,e; Supplementary Fig. 6). To determine whether more subtle defects in intrahepatic biliary transport were present, we used the phospholipase-dependent fluorescent phospholipid PED6 transport assay as previously described^34,35^. Larval zebrafish were immersed in PED6-containing water, allowing PED6 uptake through the digestive tract and processed by the phospholipase A2 (PLA2) to generate a fluorescent product that is subsequently transported through the hepatobiliary system and accumulates in the gallbladder under normal physiological conditions^34,35^ (Fig. 2f,g). LM-depleted *irf8* mutants showed robust gallbladder accumulation of fluorescent PED6 indistinguishable from controls under both normal and elevated-cholesterol diets, indicating preserved biliary transport in vivo (Fig. 2g). Thus, despite their close association with the biliary network, LMs are not required for liver growth, biliary architecture, or bulk biliary transport, prompting us to investigate their role in liver function.

We next examined the functional consequences of LM loss using whole-liver bulk RNA-seq from adult control and *irf8* mutant zebrafish, taking advantage of the abundance of LMs and accessibility of adult zebrafish liver (Supplementary Fig. 7). Whole-liver transcriptomics revealed that cholesterol biosynthesis pathway was the most strongly altered in LM-depleted livers, accompanied by changes in transporter and metabolic genes (Supplementary Fig. 7), pointing to cholesterol homeostasis as a prominent function affected by LM loss. To determine whether LMs themselves possess the molecular machinery to contribute to these processes, we FACS-isolated *mpeg1:GFP*+ LMs and non-LM hepatic cells for bulk RNA-seq. Cell identity was confirmed by strong enrichment of macrophage signature genes in LMs and negligible expression in non-LM hepatic cells (Supplementary Fig. 7). Notably, LMs expressed high levels of genes involved in cholesterol handling, including *abca1b, abcb6b, abcc3, abcc6a*, and *lipf*, as well as genes associated with immune function, including *sting1, tlr8b*, and *mhc2dgb* (Supplementary Fig. 7), supporting a direct role for LMs in hepatic cholesterol handling and immune defense.

Consistent with our previous characterization (Fig. 1bc), residual *mpeg1*+ cells in adult *irf8* mutant livers were smaller and morphologically altered and not consistently localized with biliary epithelial cells (Supplementary Fig. 7). Given the strong influence of dietary cholesterol on cholesterol homeostasis, we next examined cholesterol synthesis under controlled normal- and high-cholesterol diets. Both control and *irf8* mutant livers appropriately suppressed cholesterol synthesis in response to high dietary cholesterol; however, LM-depleted livers also showed reduced cholesterol synthesis under normal feeding (Supplementary Fig. 7). This suggests that LM loss may increase cholesterol flux to the hepatocytes under basal dietary conditions, triggering feedback suppression of endogenous cholesterol synthesis in the liver. Although bulk RNA-seq showed increased expression of cholesterol biosynthetic genes in *irf8* mutant livers, qPCR under controlled dietary conditions and with a larger sample size revealed reduced expression. The latter likely more accurately reflects the LM-depleted phenotype, as dietary conditions were not controlled in the bulk RNA-seq samples.

### Liver macrophages directly manage circulating and dietary cholesterol in vivo

Because whole-liver and LM-specific transcriptomics implicated LMs in cholesterol homeostasis, we next asked whether LMs could directly interact with circulating substances in vivo, despite their localization outside the hepatic vasculature. This question is central to understanding how LMs could regulate hepatic cholesterol homeostasis, as the liver receives cholesterol from multiple sources that converge through the circulation: dietary cholesterol delivered from the gut as chylomicron remnants, cholesterol returning from peripheral tissues in circulating lipoproteins, and cholesterol synthesized de novo within hepatocytes^1–3^.

Whole-liver time-lapse imaging in intact larval zebrafish revealed highly dynamic interactions between LMs and circulating fluorescent tracers (Fig. 3a; Supplementary Movie 6). Although resident LMs remained associated with the biliary epithelial cells, they extended dynamic processes into adjacent hepatic vessels, as well as more stable bifurcating processes that surrounded vessels and spine-like protrusions in the vessels containing fluorescent particles from the circulation. To test whether this vascular access reflected selective uptake, we performed a short-term uptake assay following intravenous injection^36^ of diverse fluorescent tracers. Strikingly, LMs preferentially accumulated cholesterol and cholesterol-containing LDL particles, whereas substantially less uptake of short-chain fatty acids (SCFAs), lipopolysaccharide (LPS), or albumin was detected (Fig. 3b–d). Thus, despite their extravascular localization, LMs are positioned to directly sample the circulation and selectively capture circulating cholesterol.

**Fig. 3.**
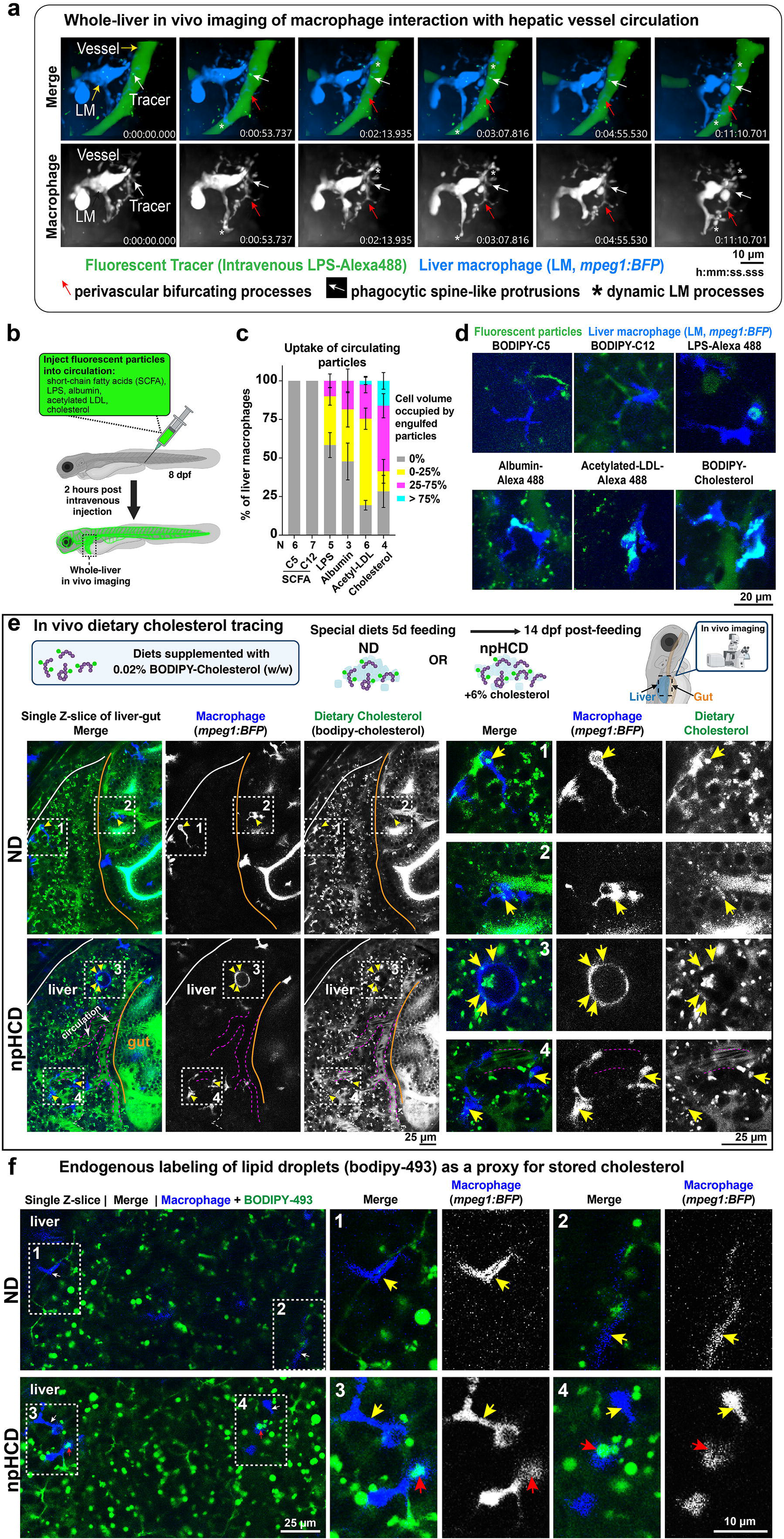
Liver-resident macrophages dynamically survey the circulation and capture circulating and dietary cholesterol. **a** High-magnification time-lapse spinning-disk confocal imaging of a single LM (BFP+) dynamically interacting with sinusoidal circulation containing LPS-Alexa488. LMs extend elaborate, highly dynamic processes, including bifurcating extensions that wrap around the vessel (red arrow) and spine-like protrusions that engulf circulating particles (white arrow). Selected cropped frames of a single LM from the time-lapse sequence corresponding to Supplementary Movie 6 are shown. **b** Schematic of the intravenous tracer uptake assay. **c** Quantification of tracer uptake by LMs, expressed as the percentage of LM volume occupied by fluorescent tracer, showing preferential accumulation of cholesterol and cholesterol-containing LDL particles. Error bars, ± SEM; N, number of animals analyzed. **d** Representative confocal images of LMs in live 14 dpf zebrafish 2 hours after intravenous injection of fluorescent tracers, showing differential tracer association with LMs. **e** Schematic of the in vivo dietary cholesterol tracing strategy and imaging region encompassing the liver and proximal intestine. Representative confocal images of control *mpeg1:BFP+* zebrafish after 5 days of feeding with BODIPY-cholesterol at 0.02% (w/w) in normal diet (ND) or non-pathological high-cholesterol diet (npHCD; supplemented with ∼6% cholesterol). White outlines delineate the liver from the periphery, and the orange line marks the liver–proximal intestine boundary. Left, tissue-level views; right, higher-magnification views of boxed regions that are numbered showing individual LMs (1, 3, 4) and one intestinal macrophage (2) containing diet-derived fluorescent cholesterol tracer (yellow arrows). Blood vessel containing circulating particles is indicated by magenta dotted lines. npHCD increases dietary cholesterol tracer accumulation in the liver and intestine. **f** Representative confocal images of live 14 dpf zebrafish livers stained with BODIPY-493 to visualize neutral lipid droplets. npHCD increases hepatic lipid droplets, whereas few LMs are associated with lipid-droplet signal.

We next asked whether LMs similarly participate in handling dietary cholesterol. We established a controlled dietary paradigm to trace fluorescently labeled cholesterol from the intestine to the liver under a normal diet (ND) or a non-pathological high-cholesterol diet (npHCD), supplemented with an additional 6% cholesterol (w/w) relative to ND, and an equal trace amount of 0.02% of the fluorescently labeled cholesterol (C24 BODIPY-cholesterol tracer)(w/w) added to both diets (Supplementary Fig. 8). Experiments shown earlier involving differential feeds also used the same dietary paradigm without the tracer (Fig. 2 and Supplementary Fig. 7). This fluorescent cholesterol tracer has previously been shown to mimic physiological functions akin to endogenous cholesterol due to its favorable location of the boron dipyrromethene difluoride (BODIPY) fluorophore modification at the very end of the side chain at C24, leaving the steroid nucleus intact enabling it to behave similarly to native cholesterol for membrane association and intracellular trafficking^37–39^. Importantly, 3-or 5-day feeding of ND or npHCD beginning at 9 dpf did not cause overt developmental or hepatic abnormalities, with control and *irf8* mutant larvae showing comparable morphology and growth across diets. Consistent with increased dietary lipid accumulation, BODIPY-493 staining, a hydrophobic dye that becomes strongly fluorescent in a nonpolar lipid-rich environment^40,41^, revealed increased neutral lipid droplets in hepatocytes and enterocytes following npHCD feeding (Supplementary Fig. 8). Neither diet induced detectable activation of resident macrophages nor changes in larval size or health (Supplementary Fig. 8). Together, these findings establish npHCD as a physiological cholesterol-loading paradigm that increases dietary cholesterol availability without overt pathology or inflammation.

Using this paradigm together with dietary cholesterol tracing, we mapped the distribution and cellular uptake of dietary cholesterol across the gut and liver (Fig. 3e). Fluorescently labeled dietary cholesterol was abundant in the intestinal lumen and enterocytes and was subsequently detected throughout the liver, with greater accumulation in hepatocytes and the hepatic circulation following npHCD (Fig. 3e). Notably, traced dietary cholesterol accumulated in hepatocytes predominantly as small punctate structures rather than the large spherical lipid droplets as seen by BODIPY-493 labeling, consistent with localization within free cholesterol-containing intracellular vesicles, rather than esterified cholesterol stored in lipid droplets (Fig. 3e,f). Importantly, fluorescent dietary cholesterol was also directly detected within LMs (Fig. 3e), providing evidence that resident macrophages capture dietary cholesterol after its delivery from the gut. In contrast, BODIPY-493 revealed a substantial increase in large endogenous lipid droplets following npHCD feeding, yet lipid droplets were only rarely observed within LMs, even under elevated cholesterol feeding (Fig. 3f). Together, these findings establish LMs as strategically positioned, highly dynamic cholesterol sensors that directly access circulating cholesterol-containing particles and capture dietary cholesterol delivered from the gut, revealing a previously unrecognized macrophage-mediated route for buffering and redistributing cholesterol within the liver.

### Single-cell transcriptomics reinforces liver macrophage cholesterol handling and reveals consequences of loss of mature LMs for hepatocyte metabolic and immune states

To determine how LMs and hepatocytes respond and adapt to physiologically elevated cholesterol by diet, and how their responses may coordinate to maintain hepatic cholesterol homeostasis, we performed whole-liver single-cell RNA sequencing (scRNA-seq) in wild-type (WT) and *irf8* mutant larvae fed either normal diet (ND) or non-pathological high-cholesterol diet (npHCD), the dietary paradigm used throughout this study (Fig. 4). At 14 dpf, following 5 days of feeding, we profiled >40,000 cells from whole livers across eight conditions, with two independent samples per condition and five whole livers pooled per sample (Fig. 4a). Most larvae carried the macrophage *mpeg1:GFP* reporter, providing an additional marker for myeloid cells. Seurat analysis of quality-filtered cells yielded ∼4,000–6,000 cells per sample and resolved 41 clusters encompassing the major hepatic cell types, including hepatocytes, biliary epithelial cells, endothelial cells, hepatic stellate cells, macrophages, and other leukocytes (Fig. 4b–d).

**Fig. 4.**
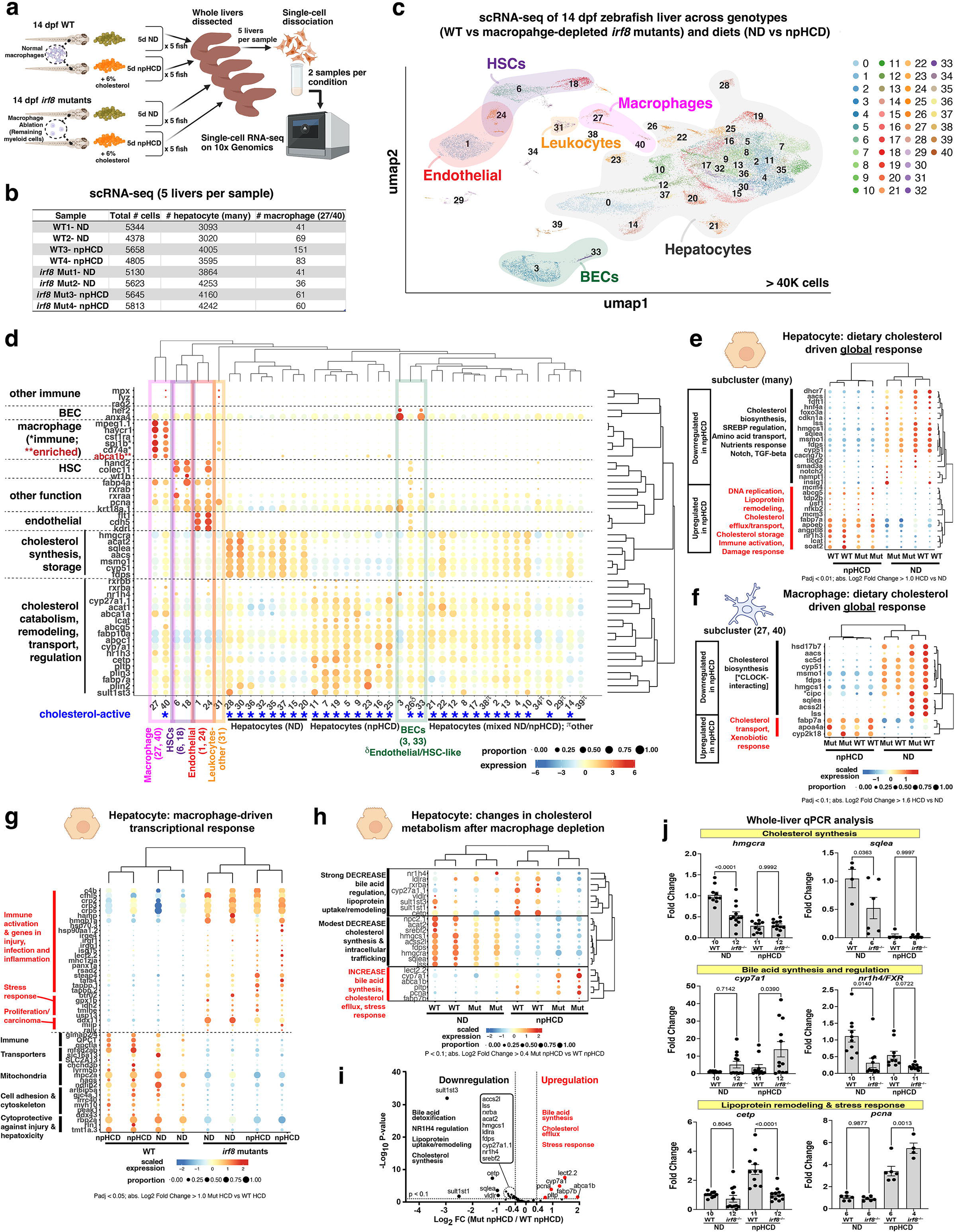
Single-cell RNA-seq reveals cell-type-specific responses to dietary cholesterol and altered hepatocyte metabolic state following liver macrophage depletion. **a** Schematic of whole-liver single-cell RNA-seq (scRNA-seq) from 14 dpf zebrafish. Two biological samples were analyzed per condition, with five pooled livers per sample. **b** Number of cells retained after quality filtering for each sample. **c** UMAP of the integrated liver scRNA-seq dataset, identifying 41 transcriptionally distinct clusters (0–40) representing major hepatic cell populations, including hepatocytes, biliary epithelial cells, endothelial cells, hepatic stellate cells, macrophages, and other leukocytes. **d** Bubble plot of canonical marker genes used to assign cell identities. Hepatocytes cluster according to dietary condition (ND or npHCD). “Cholesterol-active” denotes clusters with high expression of genes involved in cholesterol synthesis, transport, storage, or metabolism. **e,f** Bubble plots showing the most significant differentially expressed genes (DEGs) associated with dietary cholesterol in hepatocytes (**e**) and LMs (**f**). **g** Bubble plot showing the most significant DEGs in hepatocytes associated with LM depletion in *irf8* mutants compared with WT controls. **h** Bubble plot highlighting cholesterol-related genes altered in hepatocytes following LM depletion. **i** Volcano plot of DEGs between *irf8* mutant and WT hepatocytes from npHCD feeding; significantly upregulated and downregulated genes from the liver scRNA-seq dataset are shown in red and black, respectively. **j** Whole-liver qPCR validation of selected transcriptional changes identified by scRNA-seq, including genes involved in cholesterol synthesis (*hmgcra*, *sqlea*), bile acid synthesis/regulation (*cyp7a1*, *nr1h4/FXR*), lipoprotein remodeling (*cetp*), and stress/cell-cycle responses (*pcna*). Data are mean ± SEM. Statistical significance was assessed using a two-tailed Student’s *t*-test. Numbers below bars indicate animals analyzed.

Hepatocytes comprised >70% of sequenced cells and were represented by multiple clusters that separated according to the diet and genotype (73 ± 4% in WT versus 74 ± 1% in *irf8* mutants, *p* = 0.58). Based on sequenced cells, WT livers contained ∼1.8 ± 0.7% macrophages, whereas *irf8* mutants retained only 0.9 ± 0.2% residual myeloid cells (*p* = 0.05), corresponding primarily to clusters 27 and 40 (Fig. 4b). The *mpeg1+* cells in WT expressed robust macrophage identity and functional genes, including *c1qa/b/c, c1ql1, ccl34b.1, cd74a/b, csf1ra, havcr1, mfap4.2, mpeg1.1, mrc1b, mhc2, tlr3, tlr8b*, and *sting1*, whereas these mature macrophage signatures were significantly downregulated in residual *irf8* mutant myeloid cells that are characteristically *mpeg1^low^; cd68^high^; spi1b^high^* (Supplementary Fig. 9). Instead, residual mutant cells highly expressed progenitor-associated and immune-response genes, including *cd68, spi1b, pcna, s100a10b, lgals9l1, cd209l1, lect2.1, ifi44a1, irf1b*, and *irf9*, together with genes associated with lysosomal function and intracellular trafficking (*atp6ap2, ctsc, lamp2, lipf, npc2.2,* and *pikfyve*) (Fig. 4 and Supplementary Fig. 9). These molecular features matched their previously described phenotypes characterized by abnormal morphology lacking cellular processes and localization to the vasculature rather than the biliary epithelial cells (Fig. 1 and Supplementary Fig. 9). Thus, the residual myeloid cells in *irf8* mutants represent immature cells rather than functional replacement of LMs. Notably, npHCD increased the abundance of *mpeg1+* cells in both genotypes, with the strongest increase in *irf8* mutants (*p* = 0.056), suggesting that elevated dietary cholesterol stimulates myeloid production or recruitment, particularly in the progenitor-like population that persists after LM loss (Fig. 4b).

We then addressed how hepatocytes, the principal hepatic cells responsible for cholesterol metabolism, and LMs respond to increased dietary cholesterol with and without mature LMs (Fig. 4e,f). npHCD broadly suppressed expression of genes mediating cholesterol biosynthesis in both hepatocytes and macrophages independent of genotype, consistent with the known negative feedback inhibition mediated by SREBP (sterol regulatory element binding protein) to downregulate cholesterol synthesis in response to increased intracellular cholesterol^42^ (Fig. 4e,f). In hepatocytes, this response was accompanied by reduced expression of amino acid transport, nutrient-response, Notch (*notch2*), and TGF-β (*smad3a*) genes, together with increased expression of DNA replication, lipoprotein remodeling, cholesterol transport/efflux, cholesterol storage, and immune- and damage-associated genes (Fig. 4e and Supplementary Fig. 8f,g). Macrophages likewise suppressed cholesterol biosynthesis while increasing lipid transport genes, including *fabp7a*^43^ and *apoa4a*^44^, and xenobiotic-response genes such as *cyp2k18*^45^, while reducing expression of a CLOCK-interacting gene *cipc*^46^ paralog (*si:ch211-132b12.7*)(Fig. 4f). Furthermore, npHCD did not trigger an inflammatory transcriptional response^22^ in either wild-type macrophages or residual mutant myeloid cells, with *nfkb2, acod1, nfkbiaa, il1b,* and *tgfbi* remaining uninduced, consistent with a non-pathological response to elevated dietary cholesterol by npHCD as previously described (Supplementary Fig. 8g). These data reinforce the direct responsiveness of LMs to dietary cholesterol and support their active participation in hepatic cholesterol handling. They also show that dietary cholesterol elicits coordinated metabolic responses in hepatocytes and macrophages.

We next asked how loss of mature LMs alters hepatocyte state under changing diets (Fig. 4g–i and Supplementary Figs. 10, 11). *irf8* mutant hepatocytes showed coordinated induction of immune-, stress-, and cell-cycle-associated programs across diets, although the changes were intensified by npHCD. Immune genes included *c4b, cfhl5, crp* family members*, isg15, lect2.2, steap4*, and *tapbp.1/2*; stress-associated genes^25^ included *btr02, ddx11, gpx1b, hsp70.3, idh2, tmlhe*, and *usp13*; and cell-cycle-associated genes included *ddx11, miip, raly*, and *pcna* (Fig. 4g,h and Supplementary Fig. 10). UMAP analysis across all conditions further illustrate the spatial induction of immune-(*c4b, cfhl5, crp5, isg15*) and stress-(*btr02, ddx11, gpx1b, hsp70.3*) response genes^25^, in addition to *hmgb1a*^26^ linking both responses in *irf8* mutants (Supplementary Fig. 10). The induction of *isg15*^47,48^, an interferon-inducible gene, together with *hsp70.3*^49,50^ in mutant hepatocytes (Supplementary Fig. 10), suggests that LM loss may cause hepatocytes to be in a heightened immune and cellular stress state, possibly more susceptible to infectious triggers.

LM loss also shifted hepatocyte cholesterol metabolism toward cholesterol disposal. *irf8* mutant hepatocytes upregulated *cyp7a1*, which promotes bile acid synthesis, and *abca1b*, which mediates cholesterol efflux, while downregulating *hmgcra* and *sqlea* (cholesterol synthesis), *cetp* (lipoprotein remodeling), *sult1st1/sult1st3* (bile acid sulfation), and *nr1h4/FXR* (bile acid regulation)(Fig. 4g–i and Supplementary Fig. 11). Together, these changes are consistent with increased hepatocyte cholesterol burden following LM loss and compensatory engagement of cholesterol disposal pathways. Whole-liver qPCR independently confirmed reduced cholesterol synthesis genes, *nr1h4/FXR*, and *cetp*, as well as increased *cyp7a1* and *pcna* in *irf8* mutants (Fig. 4j).

Because cell-cycle-associated genes were transcriptionally elevated in residual *mpeg1+* mutant cells and hepatocytes, we assessed PCNA protein expression in vivo. *irf8* mutants showed increased PCNA+ myeloid cells irrespective of diet and a modest increase in PCNA+ hepatocytes after npHCD compared with WT (Supplementary Fig. 12). In mutant myeloid cells, *pcna* and *aurkb* upregulation accompanied a broader elevation of canonical proliferation genes (*mki67, cdk1, ccnb1, cdc20*)(Supplementary Fig. 12). In hepatocytes, by contrast, *pcna* was upregulated without significant increases in the proliferation genes and without changes in liver size or morphology, favoring an adaptive stress/replication response^51–55^ rather than overt proliferation (Supplementary Fig. 12). Together, the single-cell and bulk measurements and tissue analysis converge on a model in which mature LMs buffer dietary and circulating cholesterol and their loss shifts hepatocytes toward cholesterol disposal and metabolic adaptation while simultaneously activating immune and stress responses.

### Liver macrophage loss redirects incoming cholesterol toward hepatocytes and bile acid production

To directly test whether loss of mature LMs redistributes circulating cholesterol to other hepatic cells, we employed our previously established assay to quantify the uptake and cellular distribution of fluorescently labeled cholesterol in control and LM-depleted *irf8* mutant livers (Fig. 5). Because LMs captured substantially more cholesterol following intravenous delivery than during dietary exposure, which is predominantly taken up by hepatocytes (Fig. 3), short-term tracing of circulating cholesterol provided a sensitive approach to reveal the consequences of LM loss (Fig. 5a,b). We introduced fluorescent BODIPY-cholesterol into the circulation of mature 8 dpf larvae by intravenous injection, and performed confocal imaging of the liver in live larvae 2 hours post injection (Fig. 5a).

**Fig. 5.**
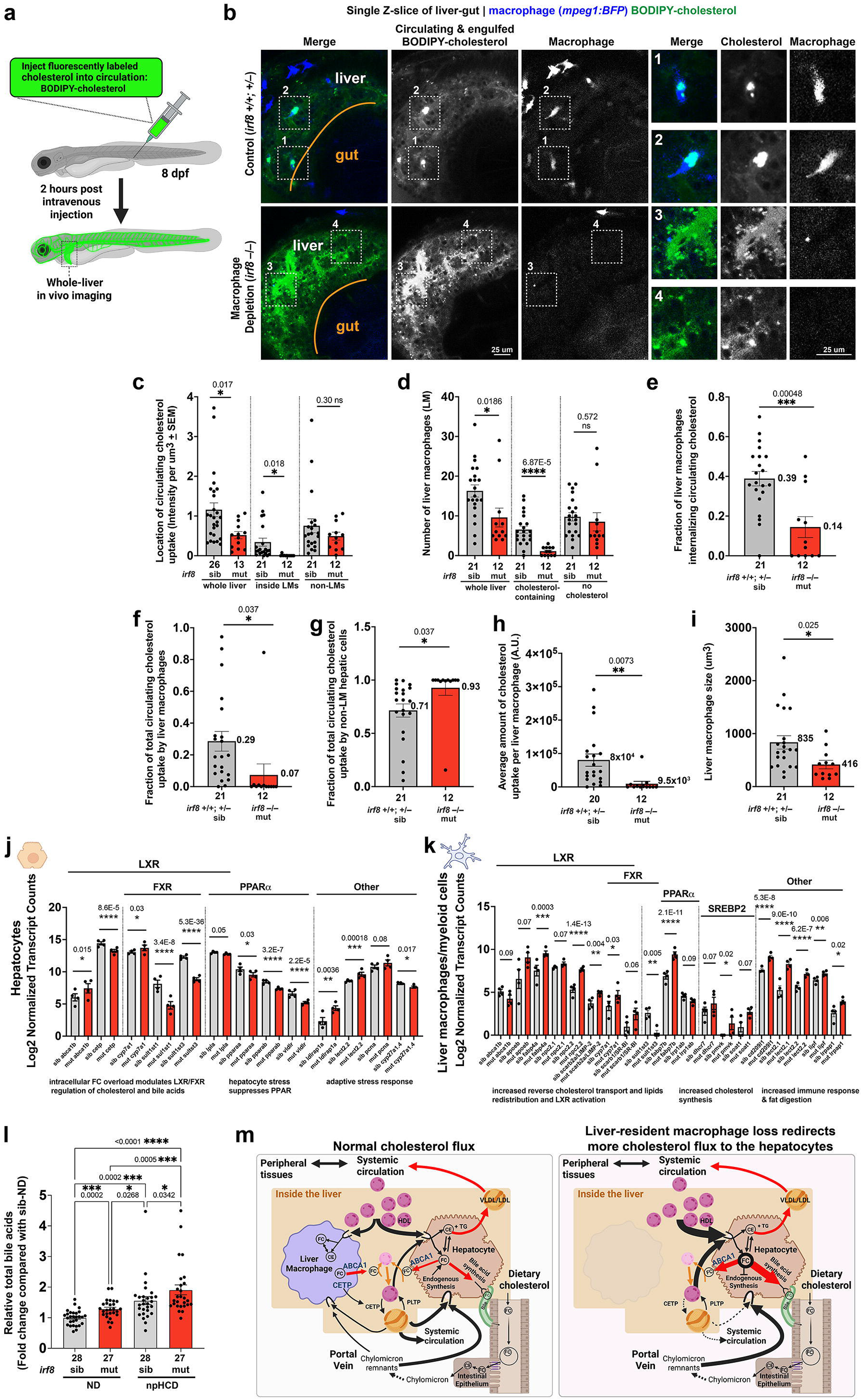
Liver macrophages buffer circulating cholesterol and promote hepatocyte adaptation and cholesterol disposal following their depletion. **a** Schematic of the intravenous cholesterol-tracer uptake assay. **b** Representative confocal images of live WT and *irf8* mutant livers 2 hours after intravenous BODIPY-cholesterol injection. WT LMs accumulate substantial cholesterol tracer, whereas *irf8* mutants lack mature LMs and show increased tracer distribution outside the LM compartment, predominantly in hepatocytes and the vasculature. Right, higher-magnification views of boxed regions that are numbered, showing tracer accumulation in LMs or residual myeloid cells. The liver–proximal intestine boundary is indicated by the orange line; circulating tracer is not detected in intestinal cells, in contrast to dietary cholesterol tracing shown in Figure 3. **c–i** Quantification of hepatic cholesterol-tracer distribution and LM uptake. Statistical significance was assessed using a two-tailed Student’s *t*-test. **c** BODIPY-cholesterol fluorescence in the whole liver, LM, and non-LM compartments. **d** Number of LMs per liver and numbers containing detectable tracer. **e** Fraction of LMs containing BODIPY-cholesterol, showing uptake by ∼40% of mature LMs compared with 14% of residual myeloid cells in *irf8* mutants. **f,g** Fraction of total hepatic BODIPY-cholesterol fluorescence localized within (**f**) or outside (**g**) the LM compartment. **h** Total BODIPY-cholesterol fluorescence per LM. **i** LM cell volume, showing reduced size of residual myeloid cells in *irf8* mutants. **j,k** Normalized scRNA-seq expression of selected genes associated with cholesterol-sensing and metabolic pathways, including LXR/FXR/PPARα/SREBP2 targets, and stress and adaptive-response programs in hepatocytes (**j**) and LMs/residual myeloid cells (**k**) from WT and *irf8* mutant livers. **l** Relative total bile acid abundance in WT and *irf8* mutant larvae fed ND or npHCD. Values from each independent experiment were normalized to the corresponding ND control. Statistical significance was determined by one-way ANOVA test followed by multiple comparisons. **m** Model proposing a role for LMs in hepatic cholesterol homeostasis. Under homeostatic conditions, LMs take up, handle, and efflux cholesterol derived from the circulation and diet, thereby buffering cholesterol exposure to hepatocytes. Following LM depletion, cholesterol is redirected toward hepatocytes, accompanied by activation of cholesterol-responsive, stress-adaptive, and immune programs and increased cholesterol disposal through bile acid synthesis. Data are presented as mean ± SEM.

**Fig. 6.**
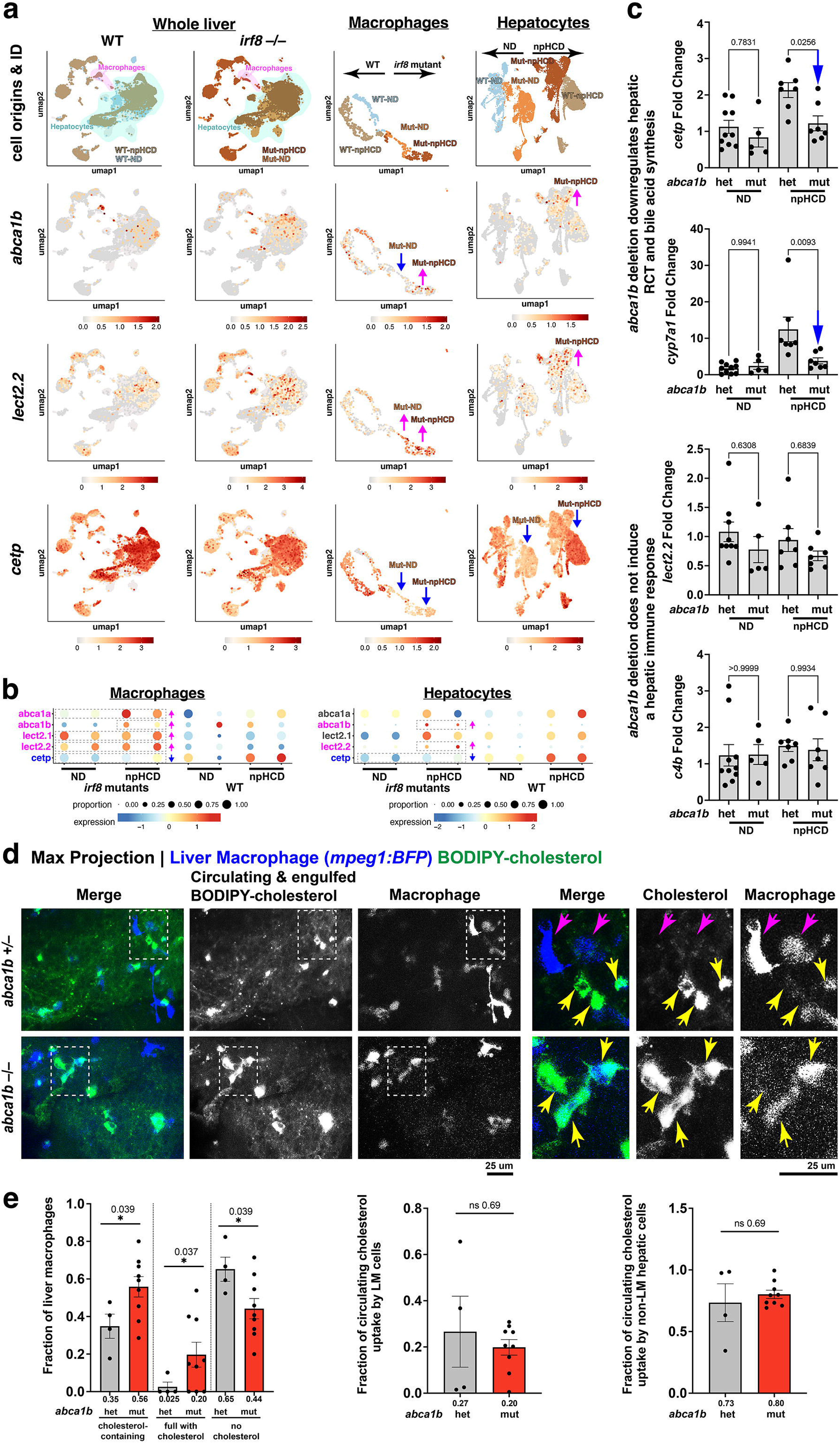
Liver macrophage depletion rewires cholesterol handling across hepatic cells, inducing abca1b and lect2.2, while suppressing cetp, while loss of macrophage abca1b downregulates hepatocyte lipoprotein remodeling and bile-acid synthesis. **a** UMAP visualization of *abca1b*, *lect2.2*, and *cetp* expression across the liver, with reclustering of myeloid cells and hepatocytes to resolve genotype-and diet-associated changes following LM depletion. Left, expression across all liver cells, with macrophage and hepatocyte populations marked by color-coded areas; middle, macrophage/myeloid-cell reclustering, showing a more macrophage-restricted expression of *abca1b* in the WT livers; right, hepatocyte reclustering across genotypes and dietary conditions. **b** Bubble plots showing the expression and proportion of cells expressing these genes and their paralogs in macrophages/myeloid cells (left) and hepatocytes (right). **c** Whole-liver qPCR analysis of *cetp*, *cyp7a1*, *lect2.2*, and *c4b* in *abca1b* mutants and control siblings under ND and npHCD conditions. **d** Representative maximum-intensity confocal projections 2 hours after intravenous cholesterol-tracer injection in *abca1b* mutants and control siblings. Left, liver tissue view; right, higher-magnification views of boxed regions showing increased accumulation of circulating cholesterol tracer within individual LMs in *abca1b* mutants. **e** Quantification of LMs with and without detectable cholesterol tracer, differentiating cells with tracer-positive versus those filled by tracer. The overall fraction of tracer-positive LMs and non-LM cells was unchanged, whereas *abca1b* mutants showed an increased fraction of LMs with tracer accumulation filling up the entire cell, consistent with impaired cholesterol efflux. N = 4 *abca1b* siblings and 9 *abca1b* mutants. Data are mean ± SEM. Statistical significance was assessed using a two-tailed Student’s *t*-test.

Quantitative cellular imaging showed that LMs accounted for nearly 30% of detectable circulating cholesterol, with ∼40% of LMs containing detectable cholesterol tracer, whereas ∼70% localized to non-LM hepatic cells, predominantly hepatocytes, in control larvae (Fig. 5b–i). In *irf8* mutants, the residual myeloid population was markedly reduced and had limited capacity to take up circulating cholesterol: only ∼14% of residual myeloid cells contained detectable cholesterol tracer, accounting for merely ∼7% of total cholesterol tracer, compared with ∼93% in non-LM cells, predominantly hepatocytes (Fig. 5b–i). Residual mutant myeloid cells were also significantly smaller, and those that internalized tracer contained significantly less cholesterol tracer than control LMs (Fig. 5h,i). Thus, loss of mature LMs directly redistributes circulating cholesterol uptake toward hepatocytes, substantially increasing the proportion of circulating cholesterol delivered to non-LM hepatic cells in *irf8* mutants.

We next asked whether this redistributed cholesterol was preferentially sequestered into neutral lipid stores. BODIPY-493 staining revealed no apparent difference in lipid droplet abundance between control and *irf8* mutant livers (Supplementary Fig. 13), despite the marked redistribution of circulating cholesterol. Together, the increased delivery of fluorescent cholesterol to hepatocytes and the absence of increased lipid droplet storage suggest that the additional cholesterol is not predominantly esterified and sequestered in lipid droplets, but instead may contribute to the intracellular free cholesterol pool. This metabolically active pool can be exported directly or converted into oxysterols and bile acids, providing routes for cholesterol disposal and engaging LXR- and FXR-associated signaling^56,57^.

Consistent with this model, hepatocyte-specific analysis of cholesterol, bile acid, and lipid-sensing (LXR, FXR, PPARα, SREBP) pathways^56–59^ from the scRNA-seq dataset revealed genotype-dependent changes in cholesterol and bile acid metabolism independent of diet. They include increased expression of the cholesterol efflux transporter *abca1b* and bile acid synthesis enzyme *cyp7a1* (Fig. 5j), whereas *cetp* was reduced, suggesting altered lipoprotein remodeling and reduced capacity for lipoprotein-mediated reverse cholesterol transport (Fig. 5j). These changes occurred alongside reduced expression of *pparaa/b*, *lpla*, and *vldlr*, indicating compromised PPARα-associated lipid metabolic capacity, while *lect2.2*, *pcna,* and *ldlrap1a* were increased, consistent with activation of stress-adaptive, proliferative, and cholesterol-acquisition programs (Fig. 5j). Thus, hepatocytes respond to the increased cholesterol delivery associated with LM loss by remodeling both cholesterol acquisition and disposal pathways, with enhanced efflux and conversion of cholesterol toward bile acid synthesis and likely altered lipoprotein-mediated cholesterol transport.

The residual myeloid cells in *irf8* mutants exhibited a distinct response at baseline under normal diet, characterized by increased expression of genes involved in intracellular lipid trafficking (*fabp4a, fabp7b, npc2.1/2, scarb2/LIMP-2*) and cholesterol uptake (*scarb1/SR-BI*), but reduced *abca1b*, suggesting increased lipid acquisition and intracellular trafficking without a corresponding capacity for cholesterol export (Fig. 5k, Supplementary Figs. 9, 11). These cells also upregulated cholesterol biosynthesis, immune-response genes (*cd209l1, lect2.1/2*), and the lipid-digestive enzyme *lipf* (Fig. 5k, Supplementary Figs. 9, 11), consistent with a metabolically and functionally altered residual myeloid state that may be poorly equipped to dispose of accumulated lipid.

Because both the cholesterol-tracing and hepatocyte transcriptional responses are consistent with the possibility of an increased cholesterol disposal through bile acid synthesis in hepatocytes in the absence of LMs, we next measured total bile acid levels in control and *irf8* mutant larvae under both diets. *irf8* mutants exhibited significantly elevated bile acid levels across diets, providing biochemical evidence that cholesterol is increasingly routed toward bile acid production following LM loss (Fig. 5l).

As an independent approach to deplete LMs, we adopted an acute clodronate-mediated macrophage depletion protocol^60^ (Supplementary Fig. 14). Although transcriptional responses were more variable, consistent with differences in the extent of LM depletion achieved by clodronate treatment, whole-liver qPCR revealed changes in the same direction as those observed in *irf8* mutants: *cyp7a1* showed a modest increase, whereas *nr1h4/FXR* and cholesterol biosynthesis genes were reduced, although these changes did not reach significance. *cetp* was significantly reduced following npHCD, while the DNA replication-associated gene *pcna* was highly variable and unchanged. Given that hepatocytes comprise most liver cells, these whole-liver qPCR measurements predominantly reflect hepatocyte transcriptional responses. Thus, despite the greater inter-animal variability and less pronounced magnitude of the response, clodronate-mediated LM depletion recapitulated the overall shift toward altered cholesterol disposal and lipoprotein remodeling observed in *irf8* mutants. The independent depletion approaches support a role for LMs in regulating hepatic cholesterol flux and disposal (Fig. 5m).

### Loss of liver macrophages disrupts cell-type-specific cholesterol-handling transcriptional program

To define mechanisms by which LMs regulate hepatic cholesterol homeostasis, we focused on *abca1b*, *lect2.2*, and *cetp* and their paralogs, selected for prominent macrophage/myeloid expression and significant changes in hepatocytes and myeloid cells following LM loss (Fig. 6). UMAP and bubble-plot analyses of these genes revealed distinct cell-type-specific expression patterns at baseline and after LM depletion in *irf8* mutants (Fig. 6a,b). RNA in situ hybridization chain reaction (HCR) further validated LM enrichment of *abca1b* and shared *cetp* expression in hepatocytes and LMs (Supplementary Fig. 15). In WT livers, increased dietary cholesterol induced *abca1a* and *cetp* in both LMs and hepatocytes, whereas *lect2.1* was selectively upregulated in hepatocytes after elevated dietary cholesterol. Notably, *abca1b* and *lect2.2* were not normally expressed by hepatocytes under either diet, but were induced in a subset of hepatocytes following LM loss. Both genes were also significantly upregulated in residual mutant myeloid cells, particularly after npHCD (Fig. 6a,b). Strikingly, LM loss disrupted these lineage-restricted programs, with *abca1b* and *lect2.2* ectopically induced in mutant hepatocytes, while both *lect2* paralogs were ectopically upregulated in residual myeloid mutant cells, particularly following npHCD (Fig. 6a,b). Conversely, *cetp* was uniformly downregulated in both cell types in *irf8* mutants despite its induction by increased dietary cholesterol in WT livers. Given the known role of *cetp* in lipoprotein remodeling and reverse cholesterol transport^61–64^, its loss suggests that LM depletion may alter not only hepatic cholesterol handling but also lipoprotein remodeling and systemic cholesterol flux. The upregulation of macrophage associated genes in mutant hepatocytes therefore suggests that LM loss alters the transcriptional response and metabolic state of neighboring hepatocytes.

### Liver macrophages buffer hepatic cholesterol flux in part by controlling its export via *abca1b,* thereby limiting cholesterol burden in hepatocytes

We next tested whether macrophage-restricted *abca1b* contributes directly to LM-mediated influence on hepatocyte cholesterol homeostasis. Using CRISPR/Cas9, we generated a stable 10-bp insertion frameshift mutation in zebrafish *abca1b*, which selectively reduced *abca1b* without affecting *abca1a* expression (Supplementary Fig. 16). In whole-liver analyses and short-term circulating cholesterol-tracing experiments, *abca1b* loss increased cholesterol tracer retained within LMs without altering tracer distribution to other hepatic cell types (Fig. 6d,e), consistent with impaired macrophage cholesterol efflux.

Despite this increased macrophage cholesterol retention, *abca1b* loss did not reproduce the immune or stress transcriptional response associated with LM depletion, as *lect2.2*, *c4b*, and *hmgb1a* were not induced. However, following npHCD, *abca1b* mutants showed reduced *cetp* and *cyp7a1a* expression (Fig. 6c), linking macrophage cholesterol export to hepatic lipoprotein remodeling and cholesterol-to-bile-acid conversion. Thus, *abca1b*-mediated cholesterol export by LMs contributes to the metabolic function of the liver and influences downstream hepatic cholesterol handling, but is insufficient to account for the dysregulated hepatocyte immune and stress states caused by LM loss.

## Discussion

Our findings support a model (Supplementary Fig. 17) in which LMs buffer hepatic cholesterol flux by capturing and exporting, in part via *abca1b*, dietary and circulating cholesterol, thereby limiting its delivery to hepatocytes while coordinating lipoprotein remodeling through *cetp* regulation and influencing bile acid synthesis for cholesterol disposal. The retention of intracellular cholesterol in *abca1b* mutant macrophages is consistent with the conserved role of ATP-binding cassette transporter A1 (ABCA1) in cholesterol efflux via ApoA1 to form nascent HDLs (Fig. 6d,e), mutations in which cause Tangier disease, characterized by severely reduced HDLs and cholesterol-laden macrophages^65,66^. Furthermore, deletion of the macrophage-restricted *abca1b* paralog reduced hepatic *cyp7a1* and *cetp* expression, showing that sequestering cholesterol in macrophages can reduce the cholesterol-processing demand on hepatocytes (Fig. 6). This further demonstrates that LMs regulate cholesterol flux to hepatocytes. Moreover, when mature LMs are lost, the cholesterol-buffering capacity of the liver is diminished, redirecting cholesterol flux toward hepatocytes and eliciting compensatory disposal and metabolic, stress, and immune-state reprogramming that is further intensified after elevated dietary cholesterol. The failure of *abca1b* loss to recapitulate the hepatocyte immune stress responses argues against impaired macrophage cholesterol efflux as the primary trigger and instead implicates loss of the upstream LM cholesterol-uptake and buffering function as a more probable driver of hepatocyte stress. In addition, aberrant induction of both *lect2* paralogs in hepatocytes and residual immature myeloid cells may enhance the altered hepatocyte state in LM-depleted *irf8* mutants. This provides a framework for understanding how disruption to macrophage ability to sense, uptake or process lipids redirects cholesterol to other cell types and may ultimately contribute to metabolic disease.

We reveal an important role for zebrafish LMs in cholesterol handling to support hepatic homeostasis. Although zebrafish LMs reside predominantly outside the vasculature rather than within hepatic sinusoids, their continuous surveillance of the circulation is functionally reminiscent of mammalian Kupffer cells^67–69^. These results raise the possibility that zebrafish LMs share features with the lipid-associated macrophage (LAM) population described around bile ducts in healthy mouse and human livers^4,5^. LAMs have largely been identified in diseased, injured or inflamed liver, including in obesity, non-alcoholic steatohepatitis, and cirrhosis, where they expand and have been shown to limit fibrosis and inflammation^5,70–72^, but their physiological function remains unclear. This study establishes cholesterol handling as a physiological function of LMs in the healthy zebrafish liver and may therefore provide insight into the homeostatic function of LAMs.

Strikingly, loss of LMs, either in *irf8* mutants or following an acute clodronate-mediated depletion, resulted in robust downregulation of *cetp,* which is normally expressed by both hepatocytes and macrophages (Fig. 6 and Supplementary Figs. 14, 15). CETP is best known for transferring cholesteryl esters from HDL to triglyceride-rich apoB-containing lipoproteins, particularly VLDL and chylomicron remnants, thereby promoting redistribution of cholesterol through the lipoprotein pool^61–63^. Its loss can therefore shift cholesterol handling toward direct hepatic uptake and disposal, including conversion to bile acids, and away from redistribution through VLDL/LDL for peripheral delivery or subsequent hepatic uptake. These findings suggest that LMs may regulate reverse cholesterol transport^73^, a process of transferring cholesterol from the periphery to the liver for excretion, not simply by removing cholesterol from the circulation, but by controlling its partitioning and processing among HDL, apoB-containing lipoproteins, and hepatocytes. Consistent with this possibility, CETP is expressed by human macrophages, where gain- and loss-of-function studies show that it promotes free cholesterol efflux^64^, and may be predominantly produced by non-hepatocyte populations^74^. However, its in vivo function remains incompletely understood. CETP may also intrinsically influence macrophage metabolic state: in CETP-expressing transgenic mice, CETP attenuates pro-inflammatory programs, limits cholesterol accumulation, and promotes an antioxidant state^75^. Notably, commonly used murine models lack functional endogenous CETP^76^, whereas zebrafish has endogenous *cetp* and thus provides a uniquely tractable vertebrate model for investigating CETP-dependent cholesterol metabolism in vivo with direct relevance to human physiology.

This study reveals both conserved and previously unrecognized features of hepatic LECT2 induction. LECT2 (leukocyte cell-derived chemotaxin 2) was originally identified as a neutrophil chemotactic factor but is now recognized as a ∼16-kDa secreted hepatokine produced predominantly by hepatocytes^27,77^. Its induction is associated with metabolic dysfunction, including obesity and insulin resistance, as well as inflammation, fibrosis, immune activation, and liver cancer^23,27,78^. Zebrafish possess three tandemly arranged *lect2* paralogs on chromosome 14, whose functions remain poorly defined but are strongly induced by infection, LPS, and live attenuated vaccination^24,79^. Of these, only *lect2.1* and *lect2.2* are expressed in the liver under different conditions (Fig. 4-6). Consistent with mammalian studies^23,78,80,81^, high cholesterol feeding significantly upregulated hepatic *lect2* in wild-type zebrafish, specifically *lect2.1* in hepatocytes (Fig. 6b). Interestingly, LM depletion in *irf8* mutants aberrantly induced both *lect2* paralogs in residual myeloid cells and *lect2.2* in hepatocytes, despite their normally negligible expression in these cells (Fig. 6b). Thus, these results raise the possibility that residual mutant myeloid cells reinforce the altered hepatocyte state through LECT2 signaling, and that LECT2 itself may mediate systemic communication during liver stress adaptation.

Macrophages share an intimate, bidirectional relationship with cholesterol, in which both the cholesterol acquired from their environment and their own intracellular cholesterol metabolism can profoundly shape their cellular state and function^82^. This relationship has been revealed largely through in vitro studies and in macrophages from adipose tissue, atherosclerotic lesions, acute dyslipidemia, and inflammatory, infectious, cancerous, and injured tissues, where changes in cholesterol availability or metabolism are closely coupled to immune responses^5,82,83^. However, it remains underexplored in the normal physiological environments in vivo. Importantly, cholesterol is not simply a metabolic substrate: its intracellular intermediates and downstream derivatives, including cholesterol precursors, oxysterols, bile acids, and steroid hormones, serve as potent signaling molecules that link lipid metabolism to macrophage function. For example, cholesterol precursors such as geranylgeranyl pyrophosphate (GGPP), desmosterol, 7-dehydrocholesterol (7-DHC), and lanosterol regulate protein prenylation, cholesterol efflux, inflammasome activity, antiviral signaling, phagocytosis, and bacterial killing^56,84^. Oxysterols such as 25-hydroxycholesterol (25-HC) and 27-hydroxycholesterol (27-HC) can restrict pathogen entry and replication while modulating inflammatory responses, macrophage polarization, and tumor-associated immunosuppression^56,58^. Additionally, bile acids signal through receptors such as farnesoid X receptor (FXR) and G protein-coupled bile acid receptor 1 (TGR5) to restrain inflammasome activation and inflammation^57,59^. Steroid hormones derived from cholesterol, including estrogens, glucocorticoids, and progesterone, likewise influence macrophage polarization, inflammatory responses, and antiviral immunity^85^. These previous studies establish the major impact of cholesterol metabolism on macrophage function, while raising a broader question: how does this metabolic dialogue operate in a tissue environment where macrophages continuously encounter and process cholesterol under normal physiology?

Our finding that resident liver macrophages are critical regulators of hepatic cholesterol homeostasis addresses this question in a physiological setting. Rather than responding to cholesterol only as a consequence of inflammation, infection, or lipid overload, liver macrophages actively participate in the normal handling and partitioning of cholesterol, integrating cholesterol acquired from the circulation with their intrinsic metabolic machinery to support hepatic and systemic lipid balance. This study reveals resident macrophages as an integral component of physiological cholesterol metabolism and expands our understanding beyond their established roles as immune sentinels to how macrophages are intrinsically equipped to sense, process, and redistribute metabolic resources within tissues. Mechanistically, we show that liver macrophages routinely import cholesterol from circulation or diet, exporting it in part using *abca1b,* and coordinate lipoprotein remodeling via *cetp,* thereby regulating cholesterol delivery to hepatocytes and elsewhere and influencing hepatic bile acid synthesis for cholesterol disposal. More broadly, defining these physiological capabilities provides a foundation for understanding the remarkable plasticity and functional breadth of macrophages, how their metabolic programs are shaped by their native environments, how these programs become maladaptive in disease, and ultimately how macrophages might be harnessed or engineered for therapeutic and regenerative applications.

## Methods

### Zebrafish transgenic and mutant lines

Wild-type, mutant, and transgenic zebrafish (*Danio rerio*) were derived from the following lines: *irf8*^st95 19^, *fabp10a:BFP*^18^, *fabp10a:dsRed*^18^, *kdrl:mCherry*^18^, *lyz:GFP*^18^, *mpeg1:GFP*^18^, *mpeg1:BFP*^22^, *krt18:GFP*^30^, *abca1b*^nc306cs^ (this paper), and *irg1-KI:GFP*^22^. Zebrafish were maintained at 28.5°C under a normal 14-hour light/10-hour dark cycle. All animal procedures were performed in accordance with protocols approved by the University of North Carolina at Chapel Hill Institutional Animal Care and Use Committee (19-132, 22-103, and 25-054). Genotyping primers and assays are described in Key Resources Table.

### Generation of *abca1b* mutant zebrafish line

CRISPR-Cas9 mutagenesis was performed as described previously^86^ to target zebrafish *abca1b* (NCBI accession XM_005173080.5; Gene ID 100136868). Briefly, multiple *abca1b*-specific guide RNAs were co-injected with Cas9 mRNA into one-cell-stage embryos to increase probability of generating loss-of-function and large-deletion alleles. T7 endonuclease assays were used to screen F0 and F1 embryos for editing. Adult F1 fish carrying CRISPR-induced mutations were selected for cloning and Sanger sequencing to identify the precise edits (Supplementary Fig. 16). Germline transmission of the 10-bp insertion frameshift allele *nc306cs* was confirmed in the F2 generation. All oligonucleotides and reagents are listed in the Key Resources Table.

### In vivo time-lapse and static confocal imaging and analysis

Imaging was performed using a Nikon A1R+ hybrid Galvano and resonant scanning confocal system equipped with an ultra-high speed A1-SHR scan head and controller, and Nikon Crest V3 spinning disk confocal system. Larvae were anesthetized with 0.02% tricaine and mounted in 1-1.5% low-melting agarose for live imaging. Z-stacks were acquired using imaging parameters optimized for each experiment, with acquisition settings kept identical between sample groups used for quantitative comparisons. Image analysis and three-dimensional reconstruction were performed using Imaris 10.1.1 (Oxford Instruments). Liver structures were segmented using the Surfaces module and biliary networks were reconstructed using the Filaments module with automated tracing followed by manual refinement. Individual liver macrophages (LMs) were subsequently segmented as three-dimensional surfaces based on *mpeg1:BFP* or *mpeg1:GFP* fluorescence. BODIPY-cholesterol signal contained within LM surfaces was compared with the total signal within the whole-liver surface to determine the proportion of detectable hepatic BODIPY-cholesterol associated with LMs. The remaining hepatic BODIPY-cholesterol signal outside LM surfaces was considered the non-LM hepatic fraction. Individual LM surfaces were additionally classified as BODIPY-cholesterol positive or - negative based on detectable GFP-channel fluorescence within each surface, and the percentage of tracer-positive LMs was calculated relative to the total number of segmented LMs within each liver. Quantitative measurements were exported for statistical analysis. For hepatocyte proliferation analysis, liver regions encompassing 15 µm were manually segmented. PCNA-positive hepatocytes and DAPI-positive nuclei were quantified using the Spots module. DAPI-positive nuclei arranged in elongated chains of five or more neighboring nuclei with ≤3 µm edge-to-edge spacing were excluded to remove endothelial nuclei.

### Dietary manipulations: ND, npHCD, and cholesterol tracing

To raise mature and developmentally normal zebrafish larvae up to 14 dpf (with a standard length [SL] of typically 6-6.5 mm), zebrafish starting by 5 dpf were maintained on L-type rotifers (*Brachionus* spp.) for 72 hours before initiation of experimental diets. Zebrafish was maintained at 28.5°C under a 14 h light/10 h dark cycle throughout their growth. For experimental diet feeding, they were fed twice daily using a micro-scoop that provides consistent amount of ∼ 2.5 mg of dry food per tank per feeding for 3 or 5 days. Golden Pearl 5–50 µm Reef & Larval Diet (Brine Shrimp Direct) as previously described^87^ was used as our base diet for preparing both the normal diet (ND) and non-pathological high-cholesterol diet (npHCD). For npHCD, cholesterol (Sigma-Aldrich, C75209) was dissolved in diethyl ether to prepare a 10% (w/w) cholesterol solution which was then used to increase the base diet by ∼6% cholesterol (w/w). ND was prepared the same way with an equivalent volume of diethyl ether without cholesterol. Diets were mixed, then dried overnight in a chemical hood to allow the diethyl ether to completely evaporate overnight at room temperature, followed by homogenization, and finally stored at −20°C to protect cholesterol integrity. Diets with fluorescent cholesterol tracer were made by supplementing ND or npHCD with equal amounts of BODIPY-cholesterol (GLPBIO, GC42964) to yield a final concentration of 0.02% or 200 µg/g diet. BODIPY-cholesterol was resuspended in DMSO and appropriate amounts were mixed into base diet, vortexed, allowed to dry overnight, stored at −20°C in the dark. See also Supplementary Figure 8 for characterization of the dietary treatments (ND/npHCD) on zebrafish larvae.

### Intravenous injections: fluorescent tracers and liposomes

All intravenous injections were conducted in zebrafish larvae anesthetized in 0.02% tricaine and immobilized dorsal-laterally in 1% low-melting agarose. Approximately 1-2 nl of a fluorescent tracer was injected into the caudal vein using pulled glass microcapillary needles, a manual micromanipulator (World Precision Instruments, M3301) and a pneumatic PicoPump PV 820 (World Precision Instruments, SYS-PV820) as previously described^36,60^. Successful injections were confirmed by visualization of the immediate distribution of the fluorescent tracer. Larvae were allowed to recover for 2 h before subsequent imaging. The following fluorescent tracers were prepared at 1 mM for BODIPY C5 (Fisher, D3834), BODIPY C12 (Fisher, D3822), BODIPY-cholesterol (GLPBIO, GC42964), and Alexa Fluor 488–conjugated albumin (Fisher, A13100), and at 1 mg/mL for Alexa Fluor 488–conjugated lipopolysaccharide (LPS) (Fisher, L23351) and acetylated low-density lipoprotein (AcLDL) (Fisher, L23380). For acute chemical macrophage depletion, ∼2 nL clodronate or PBS control liposomes (Liposoma) were injected intravenously with Alexa Fluor 568-conjugated 10-kDa dextran to verify successful injection at 10 and 12 dpf as previously described^60^. See also Supplementary Figure 14 for data using the liposomes. All reagents are listed in the Key Resources Table.

### Liver dissections

Whole livers were dissected from anesthetized larvae using 0.02% tricaine until loss of touch response at the indicated stages under a stereomicroscope. Standard length was recorded before dissection. The ventral body wall was opened along the digestive tract using fine insect dissection pins (Fisher, S24851), and each liver was separated from the gastrointestinal tissues. Dissected livers were immediately transferred to ice-cold buffer or snap-frozen in liquid nitrogen, depending on the downstream application. All dissections were completed within minutes of euthanasia to minimize RNA and protein degradation. The remaining larval tissue was collected in lysis buffer and processed for genomic DNA extraction to confirm genotype.

### Whole mount immunohistochemistry

Zebrafish larvae were fixed in 4% paraformaldehyde (PFA) for 2 h at room temperature or overnight at 4°C followed by a series of PBST washes and subsequent dehydration through a graded methanol series. Samples were rehydrated in PBST, permeabilized with proteinase K, post-fixed in 4% PFA and blocked in PBST containing 5% normal goat serum. Samples were incubated with primary antibodies overnight at 4°C followed by fluorescent secondary antibodies overnight at 4°C. Where indicated, nuclei were counterstained with DAPI. Antibodies and dilutions are provided in the Key Resources Table.

### Whole mount RNA in situ hybridization chain reaction (HCR)

8- or 14-days post fertilization (dpf) larvae were fixed overnight in freshly made 4% PFA at 4°C and stored in 100% methanol. To proceed with HCR, embryos were rehydrated, permeabilized with proteinase K, and processed through a series of reagents as per protocols previously published and provided by the manufacturer (Molecular Instruments). Probe targets and amplifiers were designed and made by Molecular Instruments and are listed in the Key Resources Table.

### Neutral lipid droplet staining

Neutral lipids were visualized in live larvae using BODIPY 493/503 (Thermo Fisher, D3922). Larvae were incubated in 1 µM BODIPY 493/503 in autoclaved system water for 1 h at room temperature in the dark, briefly washed in PBS, anesthetized, and mounted laterally in 1% low-melting agarose for confocal imaging. Identical acquisition settings were maintained within each experiment and standard length was recorded.

### PED6 hepatobiliary transport assay and analysis

Biliary transport was assessed using the fluorescent phospholipase substrate PED6 (Setareh Biotech, 7540). Larvae were fed either ND or npHCD for 3 days and assayed at 9 days post-fertilization (dpf). Larvae were incubated in 40 ml of 1 µM PED6 prepared in autoclaved system water in a 10-cm Petri dish for 1 h at room temperature in the dark. Mutant and sibling larvae were stained together and imaged without prior knowledge of genotype. Following staining, larvae were briefly rinsed in PBS, anesthetized in 0.02% tricaine, mounted laterally on their left side in 1% low-melt agarose, and immediately imaged using a Leica M165 FC fluorescence stereomicroscope equipped with a Leica DFC9000 GT camera. Imaging parameters were held constant for all larvae within each experiment. Following imaging, individual larvae were collected for genomic DNA isolation and genotyping, and genotypes were subsequently matched to the corresponding images. Gallbladder PED6 fluorescence was quantified in Fiji (ImageJ) by manually tracing the gallbladder as a region of interest and measuring its integrated fluorescence intensity and area. Mean gallbladder PED6 fluorescence intensity was calculated by dividing integrated density by region-of-interest area.

### Bulk RNA-seq sample preparations from adult whole liver or sorted macrophages

For bulk whole-liver RNA-seq samples, whole livers were dissected from adult zebrafish and dissociated into single-cell suspensions. For sorted liver macrophage bulk RNA-seq samples, whole livers were dissected from adult zebrafish carrying the macrophage reporter *mpeg1:GFP* followed by FACS sorting to collect GFP-positive macrophages and GFP-negative hepatic cells. Around 1,000-50,000 cells were collected based on GFP-positive or GFP-negative profiles using a BD Melody FACS sorter directly into RNA lysis buffer for subsequent RNA extraction. Total RNA was extracted using RNAqueous-Micro Kit following manufacturer’s protocol (Thermo Fisher, AM1931). RNA-seq libraries were generated from adult whole livers or FACS-sorted liver macrophage (LM) and non-macrophage hepatic cell populations. Three independent biological samples were analyzed per condition. Each biological sample comes from one adult liver. Low-input RNA libraries were prepared using the SMARTer HT workflow with Nextera XT and sequenced as 2 × 50-bp paired-end reads on an Illumina NovaSeq 6000 SP (whole liver) or S2 (sorted macrophages) platform. Low-input libraries were used due to the limitation of cell number from sorting liver macrophages.

### Bulk RNA-seq data analysis

Paired-end RNA-Sequencing was performed on adult whole liver tissues as well as FACS-sorted whole adult liver macrophages and non-macrophage hepatic cells. Three independent biological replicates for each condition were used for sequencing on an Illumina Novaseq6000 SP (whole liver) or Novaseq6000 S2 (sorted macrophages) using 2×50 base pairs. Library preparation consisted of the SMARTr HT+ Nextera XT for low-input samples. RNA-seq analysis followed a previously used protocol, which consists of modules and packages including Trimmomatic for data trimming, MultiQC for quality control, and BBMap for sequence alignment with the zebrafish reference genome (GRCz11). Gene expression levels were quantified using featureCounts, followed by differential expression analysis using DESeq2. Gene ontology analysis was performed on the differentially expressed genes with Metascape^88^ were conducted to understand the biological significance of differentially expressed genes.

### scRNA-seq cell preparations and sequencing

Whole livers were dissected from 14 dpf wild-type and *irf8* mutant larvae fed ND or npHCD. Two biological replicates were collected per condition, with five livers pooled per replicate. Livers were dissociated on ice in HBSS containing 1% BSA and 10 µM ROCK inhibitor. Cell concentration and viability were determined by acridine orange/propidium iodide staining using a LUNA-FX7 Dual Fluorescence Cell Counter. Single-cell libraries were generated using the 10x Genomics GEM-X OCM 3′ workflow with on-chip multiplexing. Samples were multiplexed in groups of four, and libraries were sequenced on an Illumina NovaSeq X Plus 10B flow cell (Read 1, 28 cycles; i7, 10 cycles; i5, 10 cycles; Read 2, 90 cycles).

### scRNA-seq analysis

Raw sequencing data were processed using Cell Ranger Multi with a custom GRCz11 reference containing the GFP transgene sequence. Count matrices were analyzed in R v4.4.0 and analyzed using Seurat v5.3.0. Low-quality cells were identified using miQC with a posterior probability cutoff of 0.80. Data were normalized using SCTransform, followed by principal component analysis using the first 30 principal components, UMAP dimensionality reduction and shared nearest-neighbor clustering with the selected resolution of 1.0 based on biological interpretability. Cell populations were annotated using established zebrafish liver markers and cell-type-specific module scores. Hepatocytes and macrophages/myeloid cells were reclustered using the same normalization, dimensionality reduction, and clustering workflow described above. For differential expression, raw counts were aggregated by biological replicate using Seurat Aggregate Expression and analyzed using a pseudobulk DESeq2 approach. Genes with ≥10 counts in at least two samples were retained, and samples containing fewer than five cells in the population of interest were excluded. Differential expression was modeled using experimental group (∼group), with size factors estimated using the poscounts method. Genes with adjusted *P* < 0.1 or better and |log∼2∼ fold change| > 1 were considered differentially expressed. To confirm paralog-specific detection of zebrafish *lect2* transcripts, we verified that *lect2.1*, *lect2.2*, and *lect2.3* were represented as distinct features in the Cell Ranger reference and examined read-level genomic and transcript assignments, splice-aware coverage, and sequence discrimination among the three tandem paralogs. Across all eight libraries, 17,089 reads were assigned to *lect2.2*, ∼99.9% had MAPQ 255 and NH=1, and direct sequence comparison of 17,046 reads showed that 94.7% had *lect2.2* as the best paralog match, with median mismatches of 0, 17, and 33 to *lect2.2*, *lect2.1*, and *lect2.3*, respectively, providing strong support for reliable gene-level discrimination of *lect2.2*. All scripts were executed on the UNC Chapel Hill’s Longleaf computing cluster, and outputs were organized into directories for reference creation, quality control, clustering, visualization, and differential expression. Data visualization and downstream analyses were performed using Seurat, ggplot2, and ShinyApp.

### RNA isolation and qPCR analysis

Total RNA was isolated from whole larvae, dissected larval livers or adult livers using the RNAqueous-Micro Kit (Thermo Fisher Scientific, AM1931). Depending on the experiment, biological replicates consisted of individual samples or pools of two or four larvae or livers. cDNA was synthesized from 30–200 ng total RNA using SuperScript IV Reverse Transcriptase (Thermo Fisher Scientific); the same quantity was used for each experimental dataset. qPCR was performed using PowerUp SYBR Green Master Mix on a QuantStudio 3 Real-Time PCR System in 10 µl reactions containing 0.5 µM each primer. All reactions were performed with three technical replicates. Relative expression was calculated using the ΔΔCt method and normalized to *ef1a*. All plots show data points representing biological replicates. Primer sequences are provided in the Key Resources Table.

### Total bile acid assay

Total bile acids were measured in size-matched 14 dpf larvae using the Diazyme Total Bile Acids Assay Kit (DZ042A-K). Individual larvae were homogenized in 100 µl ice-cold 0.9% saline, spun down at 14,000 rpm for 15 min at 4°C, and supernatant was transferred to a new tube. Total bile acid concentrations were determined from a standard curve according to the manufacturer’s assay chemistry and measured at 405 nm using a CLARIOstar Plus microplate reader. Samples were analyzed in technical duplicate.

### Statistical analysis

Statistical analyses were performed using GraphPad Prism 10 unless otherwise indicated. Data are presented as mean ± SEM and/or individual biological replicates. Pairwise comparisons were performed using two-tailed unpaired Student’s *t*-tests, with Welch’s correction when unequal variance was detected. Comparisons among three or more groups were performed using one-way ANOVA followed by pairwise comparisons with false-discovery-rate correction. Exact statistical tests, biological replicate numbers and significance thresholds are provided in the corresponding figure legends. Experiments were independently replicated at least twice.

## Data Availability

Data generated in this study are provided in the publication, Supplementary Information and Source Data File. RNA sequencing data is freely available at the GEO repository as GSE347162 for bulk RNA-seq, and GSE347242 for scRNA-seq.

## Acknowledgements

Many thanks to Jamie Winshell for generously sharing her technical expertise and materials, to the Matt Redinbo lab for kindly providing access to their plate reader, and to Michael Pack for the *krt18:GFP* line. We are grateful to the UNC High-Throughput Sequencing Facility staff (supported by P30-CA016086 and P30-ES010126) for library preparations and RNA-sequencing, Gabrielle Cannon of UNC Advanced Analytics Core (UNC CGIBD, P30 DK034987) for scRNA-seq library preparations, researchers of UNC Bioinformatics and Analytics Research Collaborative (BARC): Tara Brennan, Matt Niederhuber, and Austin Hepperla for consultation on scRNA-seq data analysis, and Michelle Altemara and staff of UNC Zebrafish Aquaculture Core Facility for zebrafish housing and care. This work was supported by NIH NIGMS grant R35GM124719 and Chan Zuckerberg Initiative (CZI) Phase 2 grant to C.E.S.

## Author Contributions

MH: experimental designs, data analysis and visualization, experimentation on all aspects of the study, writing; SMM and MC: experimentation and technical support; CES: conceptualization, experimental designs, data analysis and visualization, supervision, funding acquisition, writing- original draft and all revisions. All authors reviewed and edited the manuscript.

