## Supplementary material for "Resident liver macrophages buffer dietary and circulating cholesterol to maintain hepatocyte homeostasis": Hamilton_Shiau_biorxiv_supplementary_LM paper.pdf

**This file includes:**
**Supplementary Figures and Legends 1-17.**
**Supplementary Movie Legends.**

**Supplementary Figure 1.**

**a**

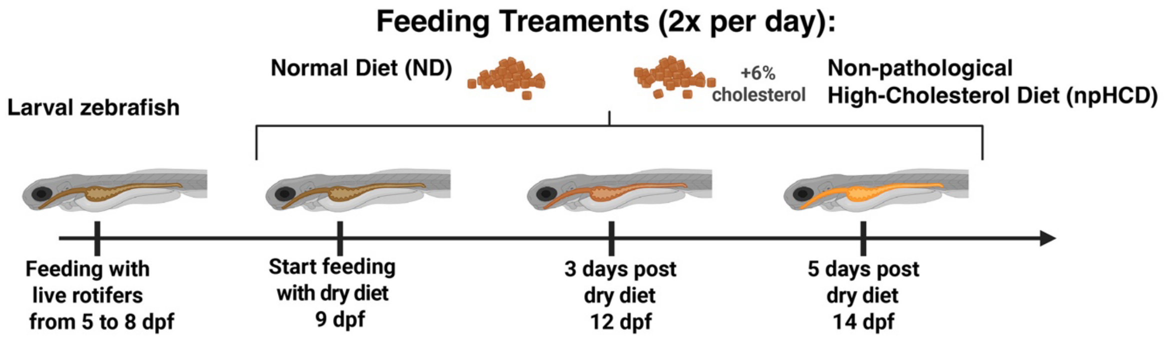


**b** **Raised in 1L tanks to support zebrafish maturation and vertical swimming, under controlled light:dark cycle (14/10 h) and temperature (28–29°C) conditions**

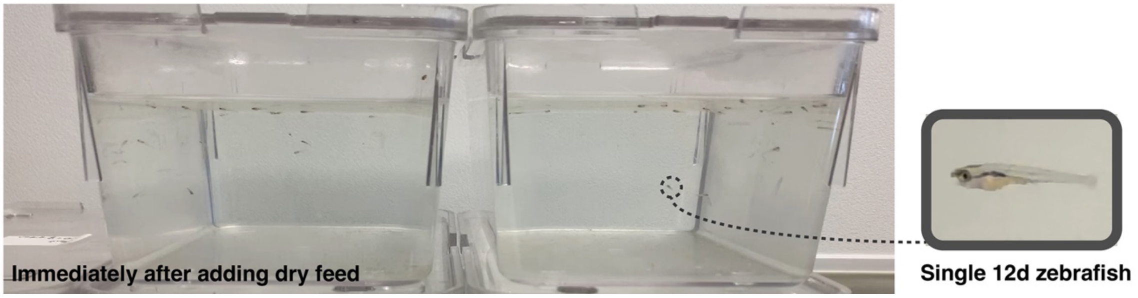


**Supplementary Figure 1 | Controlled 1-L tank rearing of larval zebrafish using live rotifer and experimental dry-diet feeding allows normal post-embryonic growth and maturation.**

**a** Schematic of the experimental feeding regimen, showing the transition from live rotifers to dry experimental diets (normal diet [ND] or non-pathological high-cholesterol diet [npHCD]). Experimental feeding was performed for either 3 or 5 days, with the 5-day regimen used primarily in this study. Live rotifer feeding for 72 hours beginning at 5 dpf to support normal digestive maturation, feeding behavior, and subsequent larval growth. Without this initial live feeding period, larval growth was markedly impaired and larvae failed to attain the size achieved after 3–5 days of dry-diet feeding.

**b** Representative image of the 1-L tank system used to rear larvae during dietary treatment. Larvae were maintained at a minimum water volume of 20 mL per larva to support normal post-embryonic growth as previously described[89,90]. The healthy growth and behavior of larvae maintained in these conditions are also shown in the corresponding Supplementary Movie 1.

**Supplementary Figure 2.**

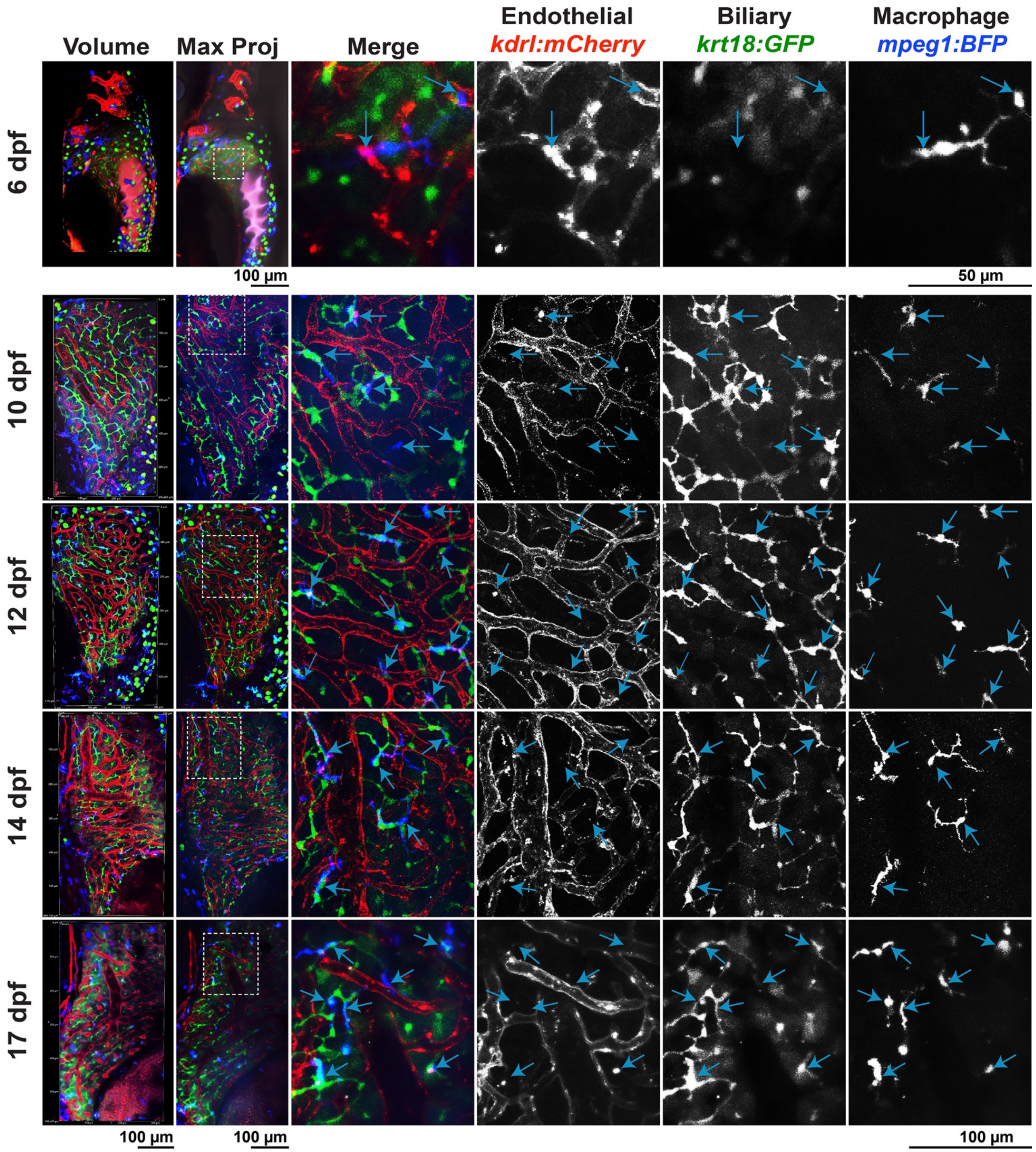

**Supplementary Figure 2 | Liver-resident macrophages localize to the biliary network while their processes can interact with vasculature during development.**

Representative 3D volume and max projection confocal images of triple transgenic zebrafish livers showing the overlay of macrophages (green), endothelial cells (red), and biliary epithelial cells (green) at 6, 10, 12, 14, and 17 days post-fertilization (dpf). High magnification of dotted box region is shown in the right panels displaying the merge and individual channels to clearly show the alignment of liver macrophages (blue arrows) with the biliary epithelial cells, but not the vasculature, although their processes can interact with the vasculature. A corresponding z-stack of the WT liver at 12 dpf is presented as a fly-through in Supplementary Movie 2 to further demonstrate how LMs are positioned relative to the biliary and vascular networks.

**Supplementary Figure 3.**

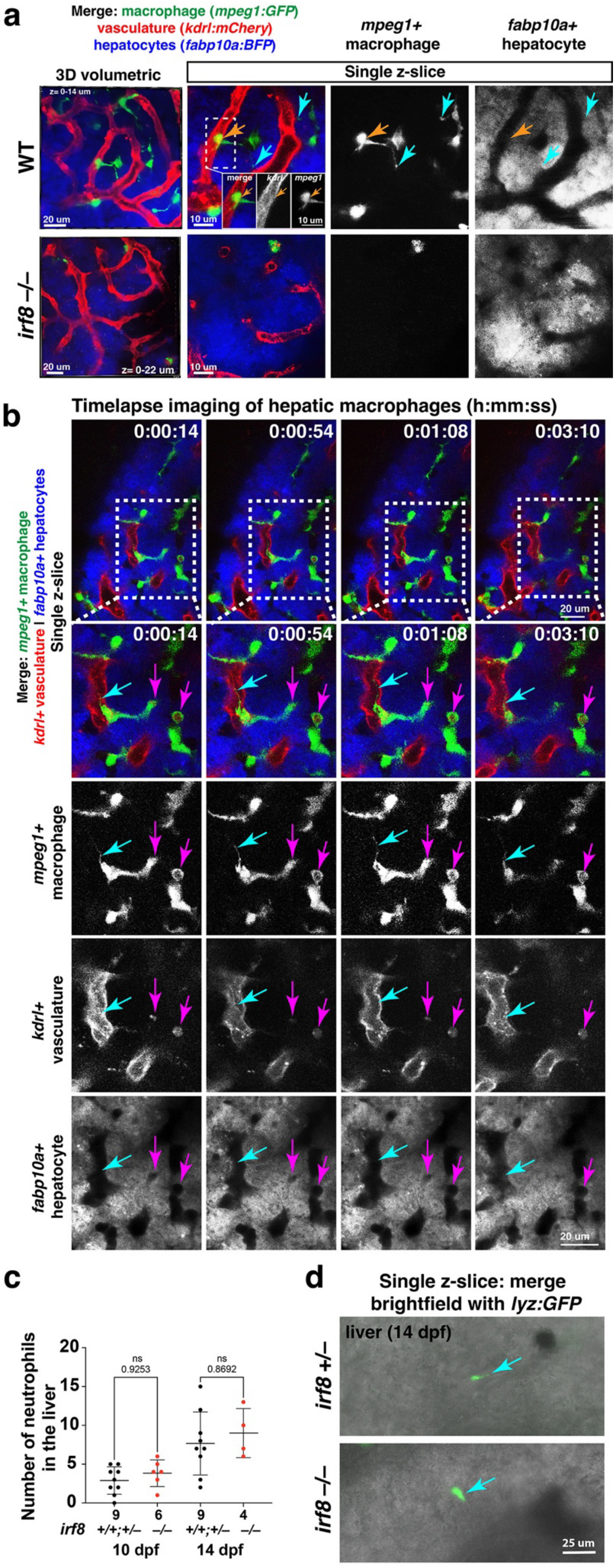

**Supplementary Figure 3 | Liver-resident macrophages are biliary-associated but dynamically sample the hepatic vasculature in zebrafish, and are lacking in *irf8* mutant livers, which contain a typical neutrophil composition comparable to controls.**

**a** Representative 3D volume view (left) and single z-slices (right) from whole-liver confocal imaging of live WT and *irf8* mutant zebrafish at 14 dpf after 5-day of normal diet. Single-channel and merged images reveal the spatial relationships among liver macrophages (LMs; green), hepatocytes (blue), and vasculature (red), highlighting LM morphology within the surrounding hepatic architecture. LM cell bodies remain closely associated with the biliary network and are largely positioned outside and away from the vasculature. However, LMs extend fine processes that frequently contact the vasculature at multiple points (blue arrows) and can partially extend across the sinusoidal endothelium into the vascular lumen (orange arrow; inset, higher-magnification view of the dotted boxed region). In contrast, residual mutant myeloid cells lack these fine processes, are smaller, and are predominantly localized around the vasculature. A corresponding movie of this liver in 3D is presented in Supplementary Movie 3 to further demonstrate how LMs are positioned and can partially penetrate through the vasculature.
**b** Representative frames from time-lapse confocal imaging show that LMs are not situated inside the endothelial sinusoids but can contain vascular-derived cargo (pink arrows) and dynamically extend and retract fine processes into the sinusoidal lumen (blue arrow) in the 8 dpf zebrafish. These dynamic extensions provide direct evidence that LMs access and survey circulation despite being predominantly localized to the biliary network. Corresponding time-lapse imaging is provided in Supplementary Movie 4. **c** Number of neutrophils in the liver using the *lyz:GFP* reporter was quantified in *irf8* mutants and sibling controls at 10 dpf and 14 dpf. There was no significant difference between the two genotype groups at both stages. Error bars show standard deviation; ns, not significant; N, number of animals analyzed is shown below each graph. **d** Representative in vivo confocal imaging of a single z-slice through the 14 dpf liver shows an example of a neutrophil observed. Brightfield was used to identify the liver tissue.

**Supplementary Figure 4.**

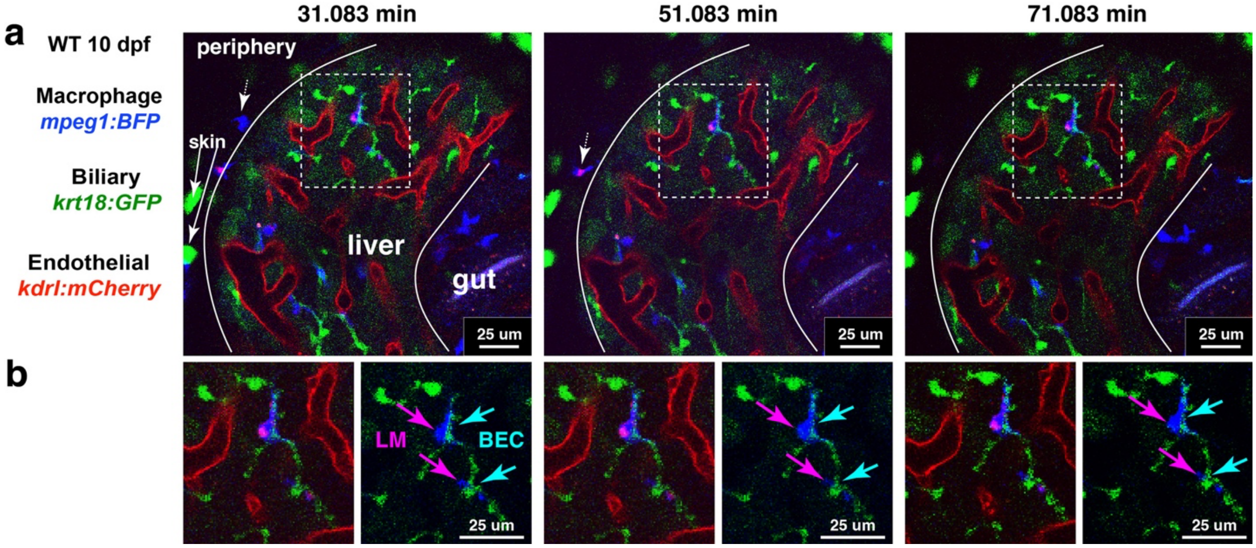


**Supplementary Figure 4 | Time-lapse imaging reveals stable positioning of liver-resident macrophages along the biliary cellular network.**

**a** Selected frames from time-lapse confocal imaging of a liver substack in a live 10 dpf zebrafish showing the stable positioning of liver macrophages (LMs; blue), whose cell bodies are juxtaposed with biliary epithelial cells (BECs; green) and remain stationary, in contrast to rapidly migrating peripheral macrophages outside the liver (white dotted arrow). LMs are not positioned along or within the vasculature. The biliary *krt18:GFP* reporter also marks a subset of skin cells (white arrows), shown as superficial rounded spots as previously described[30].

**b** High-magnification views of the boxed regions shown below each frame. Left, merged images of LMs (blue), BECs (green), and endothelial vasculature (red); right, LMs (magenta arrow) and BECs (blue arrow) overlaid to highlight their intimate alignment and contact away from the vasculature. Corresponding time-lapse imaging is provided in Supplementary Movie 5.

**Supplementary Figure 5.**

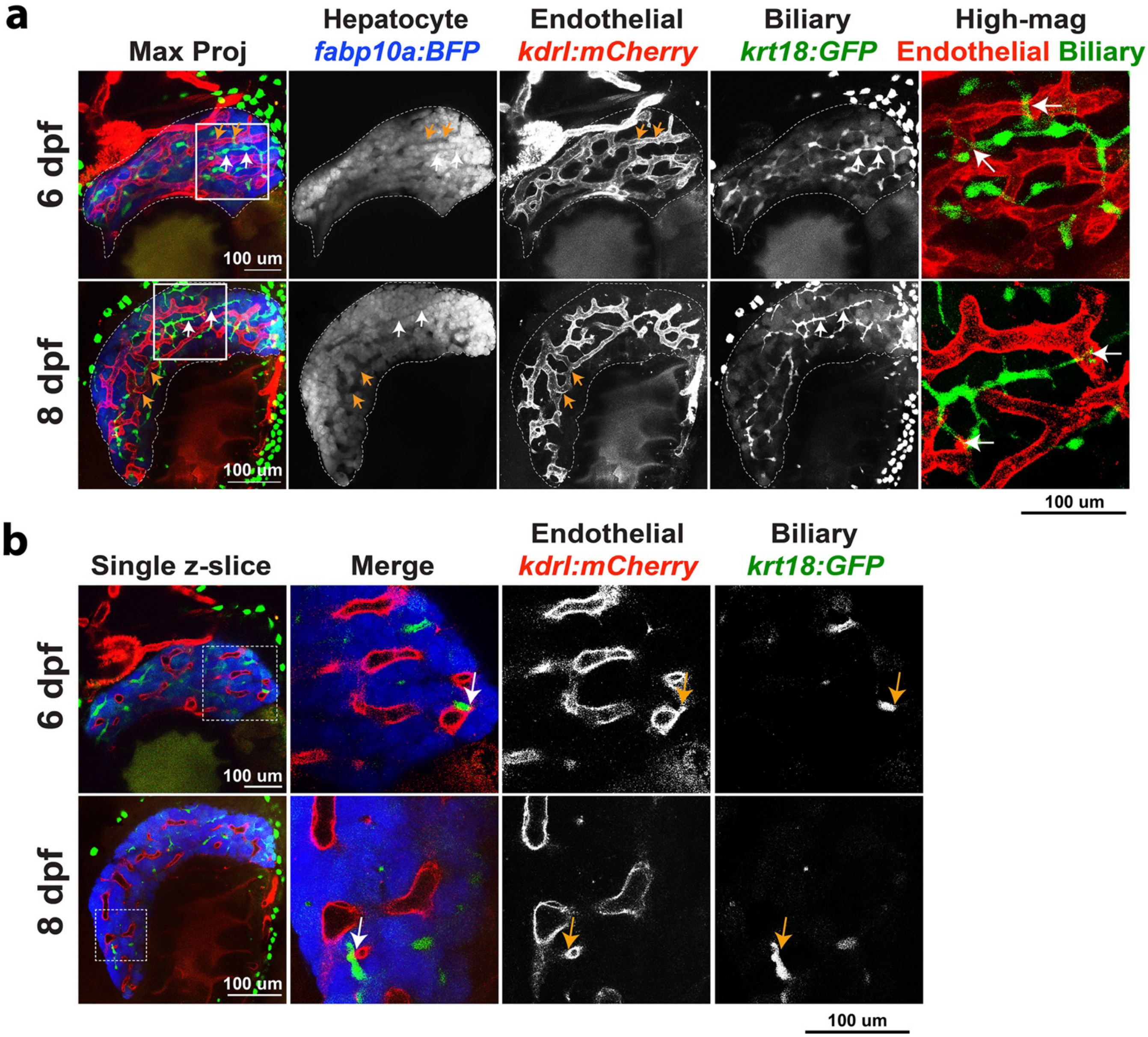

**Supplementary Figure 5 | The biliary and vascular networks form distinct, spatially organized networks with discrete points of contact.**

**a** Representative maximum projections of confocal z-stacks from whole livers of live zebrafish at 6 and 8 dpf, showing the spatial organization of hepatocytes (blue), vasculature (red), and biliary epithelial cells (BECs; green). Throughout the liver, vasculature occupies the larger interhepatocyte spaces (orange arrows), whereas the biliary network occupies thinly narrow interhepatocyte spaces (white arrows). High-magnification views of the dotted boxed regions reveal sites where the biliary network comes in contact with the hepatic sinusoidal network (white arrows). **b** Representative single z-slices corresponding to the whole-liver views in (**a**) reveal discrete sites of direct contact between BECs and endothelial sinusoids (arrows). High-magnification views of the boxed region are shown in the right panels.

**Supplementary Figure 6.**

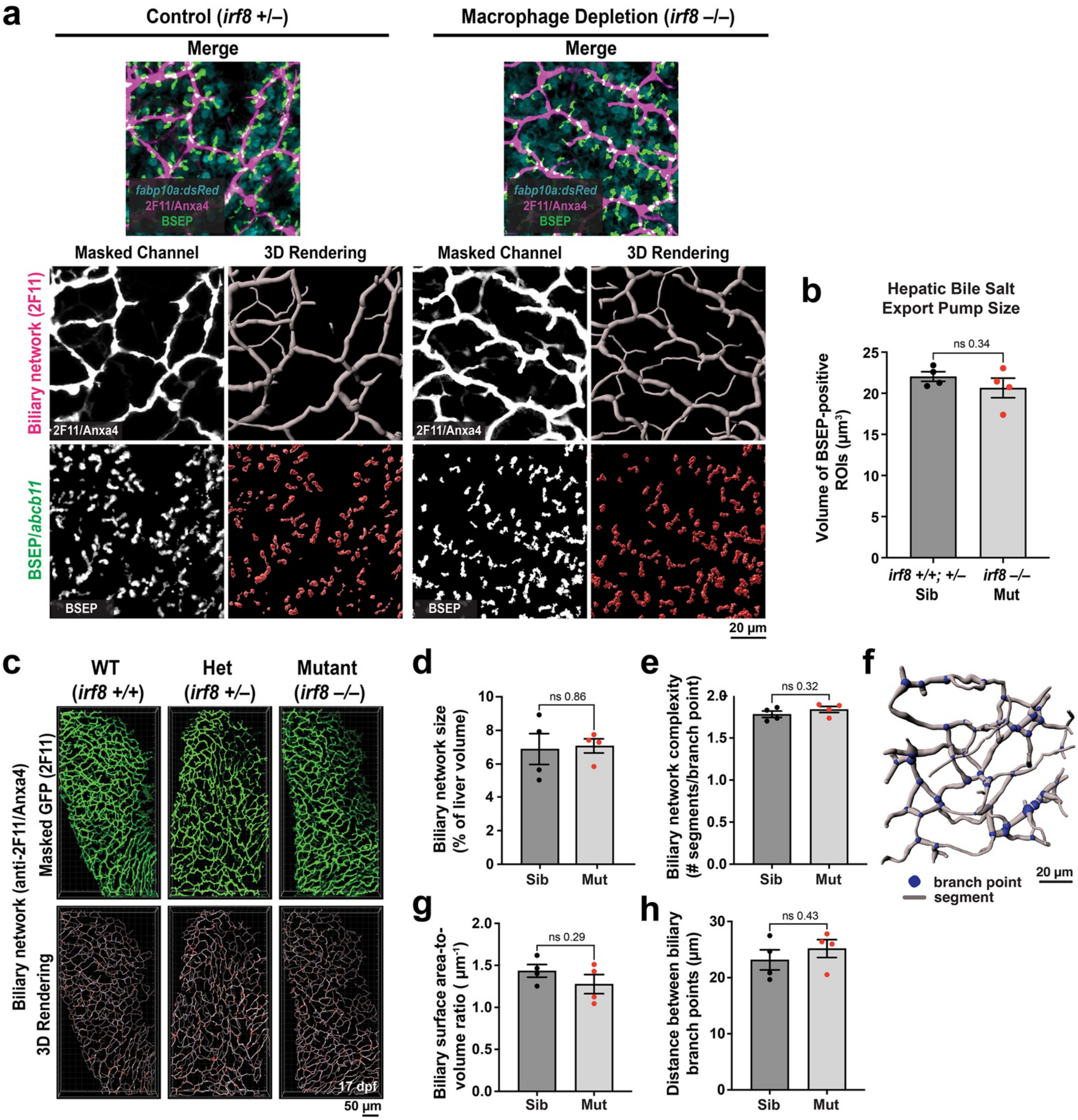

**Supplementary Figure 6 | Quantitative 3D analysis reveals preserved biliary network architecture following macrophage depletion.**

**a** Top, representative maximum projections of confocal z-stacks showing hepatocytes (blue), biliary epithelial cells (BECs; magenta), and bile salt export pump (BSEP; green), which labels the apical canalicular membrane of hepatocytes bordering the biliary epithelial cells. Images are also shown in Figure 2d. BECs were visualized by anti-2F11 immunostaining against Anxa4. Bottom, corresponding masked channels and 3D reconstructions generated using the Imaris Filament and Surface tools to quantify BSEP size and distribution, and biliary network size, complexity, and morphology in mature 17 dpf larvae.

**b** Quantification of the volume of BSEP-positive structures, representing the apical canalicular membrane compartment of hepatocytes, reveals no difference in biliary canalicular size between *irf8* mutants and sibling controls.

**c** Representative masked channel and 3D reconstruction of the entire liver biliary network based on 2F11 staining at 17 dpf. The Imaris Filament tool was used to model the biliary network.

**d–e** Quantification of biliary network volume as a percentage of liver volume and network complexity, defined as the number of biliary segments per branch point, respectively. Neither parameter differed between *irf8* mutants and sibling controls.

**f** Schematic illustrating 3D reconstruction of a biliary network. Branch points are shown as blue spheres and segments as gray bars used for quantitative analysis.

**g–h** Quantification of biliary surface area-to-volume ratio and distance between branch points, respectively. Neither parameter differed between *irf8* mutants and sibling controls. Data are mean ± SEM; each point represents an individual liver. N = 4 per group. Statistical significance was assessed using a two-tailed Student's *t*-test.

**Supplementary Figure 7.**

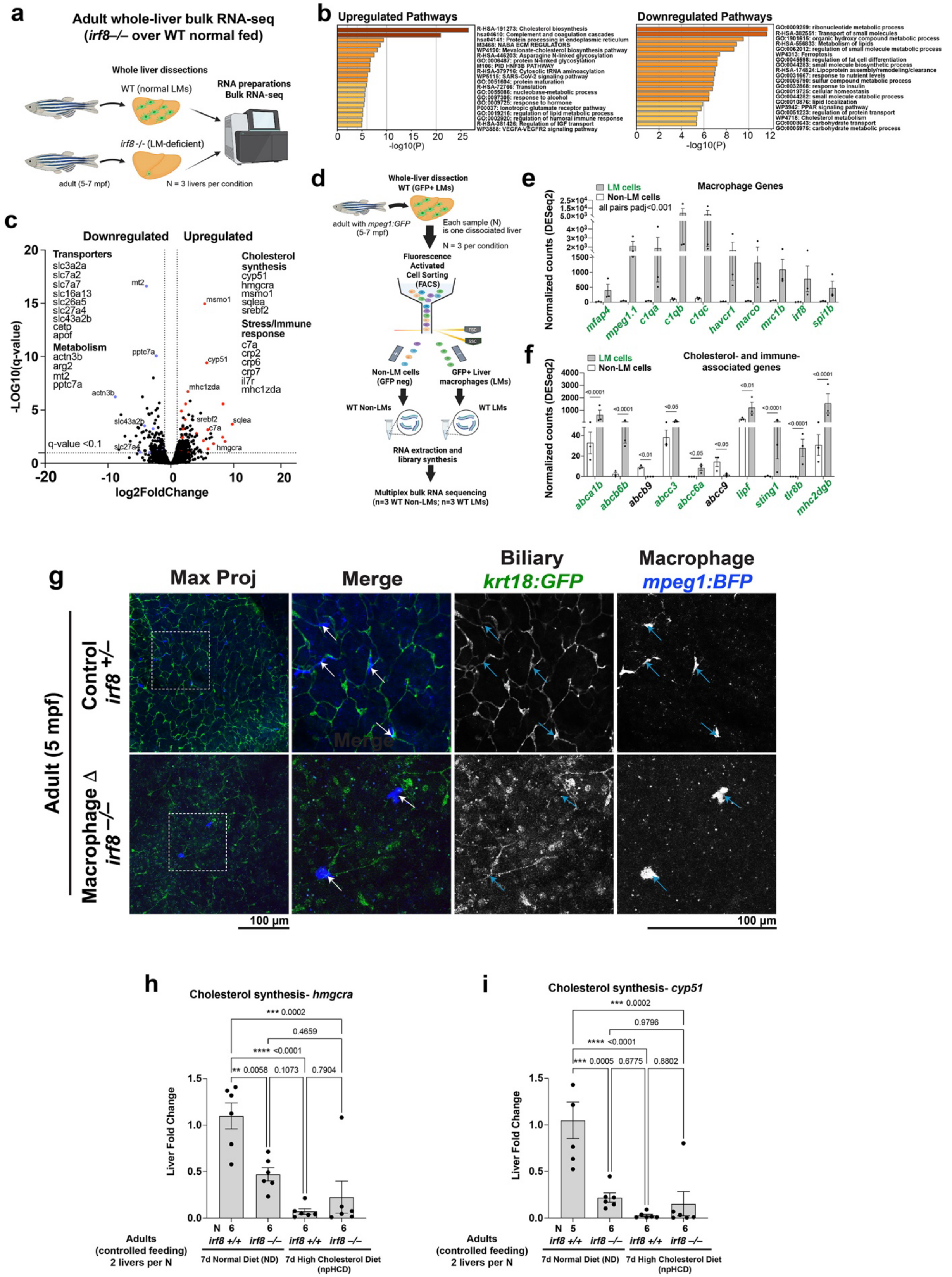

**Supplementary Figure 7 | Bulk transcriptomics identifies cholesterol metabolism as the top-ranked most altered pathway in macrophage-depleted adult livers, while adult liver macrophages express cholesterol-handling machinery.**

**a** Experimental design for bulk RNA sequencing of adult whole livers from WT and *irf8* mutant zebrafish.
**b** Enriched biological pathways identified by Metascape from differentially expressed genes (DEGs; adjusted $p < 0.05$) in *irf8* mutant versus WT livers.
**c** Volcano plot showing differential gene expression between *irf8* mutant and WT adult livers. Genes with $q < 0.1$ and $|\log_2$ fold change$| > 1$ were considered significantly changed. Selected significantly upregulated and downregulated genes are indicated.
**d** Experimental design for isolation and transcriptional profiling of adult liver macrophages (LMs). Livers from WT *mpeg1:GFP*+ zebrafish were dissociated, followed by fluorescence-activated cell sorting (FACS) of GFP-positive LMs and GFP-negative non-LM hepatic cells for bulk RNA sequencing.
**e** DESeq2-normalized expression of macrophage-associated genes in sorted GFP-positive LMs and GFP-negative hepatic cells. The strong enrichment of macrophage markers in GFP-positive cells and their minimal expression in GFP-negative cells validates the macrophage identity of the isolated GFP+ cells.
**f** DESeq2-normalized expression of selected cholesterol- and immune-associated genes in sorted LMs and GFP-negative non-LM hepatic cells. LMs showed prominent expression of cholesterol-handling genes, including *abca1b* and *abcb6b*, alongside immune-associated genes such as *sting1*, *tlr8b*, and *mhc2dgb*.
**g** Confocal imaging of live isolated adult livers from WT and *irf8* mutant zebrafish expressing *mpeg1* (macrophage) and *krt18* (biliary) reporters. LMs remain closely aligned with the biliary network into adulthood, whereas the few residual *mpeg1*+ cells in *irf8* mutants remain abnormally rounded lacking processes.
**h–i** qPCR analysis of the cholesterol synthesis genes *hmgcra* (**h**) and *cyp51* (**i**) in adult WT and *irf8* mutant livers following normal diet (ND) or non-pathological high-cholesterol diet (npHCD) feeding. Macrophage depletion was associated with significantly reduced expression of cholesterol biosynthesis genes under basal ND conditions, suggesting disruption of liver cholesterol metabolism. N, number of individual livers analyzed; mpf, months post-fertilization. Data are mean ± SEM.

**Supplementary Figure 8.**

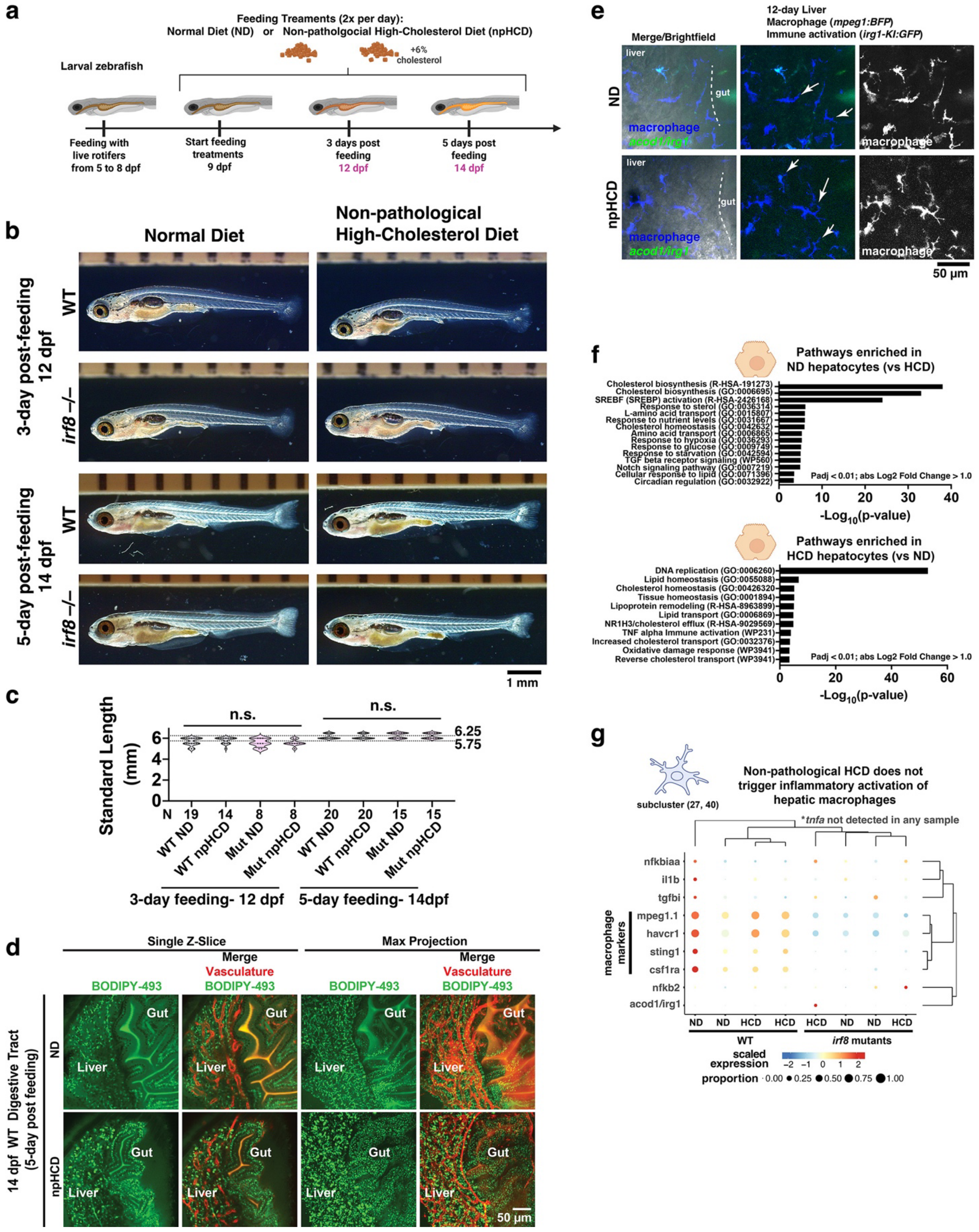

**Supplementary Figure 8 | Controlled high-cholesterol diet elevates hepatic lipid storage without growth defects or inflammatory activation, consistent with healthy larvae without overt pathology.**

**a** Schematic of the controlled dietary paradigm comparing normal diet (ND) with non-pathological high-cholesterol diet (npHCD), consisting of live rotifer feeding during early larval development followed by defined laboratory dry diets.
**b** Representative live images of WT and *irf8* mutant larvae following 3 or 5 days of ND or npHCD feeding starting at 9 dpf (grown to 12 or 14 dpf, respectively). Standard length (SL), measured from the tip of the mouth to the end of the caudal muscle, was comparable between genotypes within each feeding duration. As expected, larvae were larger after 5 versus 3 days of feeding.
**c** Quantification of SL in WT and *irf8* mutant larvae after 3 or 5 days of dietary treatment. Data are mean ± SEM; each point represents an individual larva. No significant differences were detected between ND and npHCD within either feeding duration, indicating no overt effect of npHCD on larval growth. 3-day fed zebrafish at 12 dpf had average SL size of 5.6 mm with a typical range of 5-6 mm. 5-day fed zebrafish at 14 dpf had average SL size of 6.2 mm with a typical range of 6-6.5 mm with few exceptions.
**d** Representative single z-slices and maximum projections of the liver and proximal intestine from WT larvae at 14 dpf following 5 days of ND or npHCD feeding. BODIPY-493 staining revealed increased neutral lipid droplet accumulation following npHCD feeding.
**e** Assessment of inflammatory or immune activation using an *irg1/acod1* GFP knock-in reporter (*irg1-KI*)[22] in live *irg1-KI; mpeg1* larvae at 12 dpf following 3 days of ND or npHCD feeding. Representative confocal images show no detectable induction of the *irg1* reporter in the liver, including *mpeg1*+ macrophages, following npHCD feeding.
**f** Metascape pathway enrichment analysis of liver hepatocyte transcriptomes comparing ND and npHCD conditions identifies the principal dietary transcriptional responses. The data source comes from the liver scRNA-seq dataset presented in Figure 4.
**g** Bubble plot from liver scRNA-seq showing expression of inflammatory/activation-associated genes (*nfkbiaa*, *il1b*, *tgfbi*, *nfkb2*, and *irg1*) and selected macrophage markers (*mpeg1.1*, *havcr1*, *sting1*, and *csf1ra*) across liver macrophage subclusters. Inflammatory/activation-associated genes showed no detectable diet-dependent induction, whereas canonical macrophage markers were enriched in WT LMs relative to *irf8* mutant myeloid-like populations. Together, these findings establish npHCD as a controlled cholesterol-elevating paradigm.

### Supplementary Figure 9.

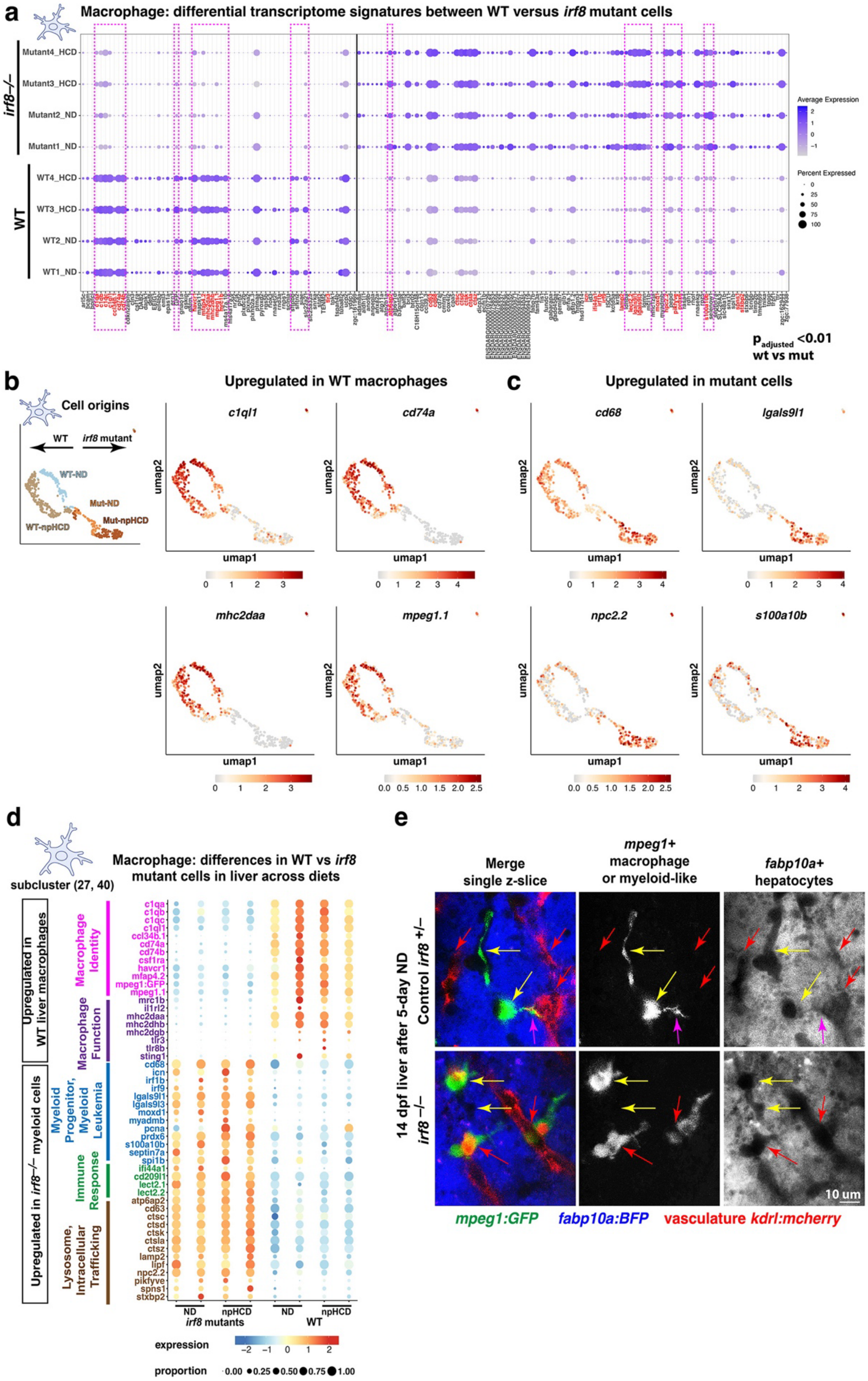

**Supplemental Figure 9 | Residual *irf8* mutant myeloid cells resemble immature cells that diverge from mature liver macrophages in identity, morphology, and localization.**

**a** Bubble plot of differentially expressed genes (DEGs; adjusted $p < 0.01$) following reclustering of macrophage/myeloid cells from the liver scRNA-seq dataset (Figure 4). WT and *irf8* mutant cells segregated transcriptionally independent of diet (ND or npHCD). WT LMs robustly expressed mature macrophage-associated genes, including *c1qa, c1qb, c1qc, c1ql1, ccl34b.1, cd74a, cd74b, csf1ra, havcr1, mfap4.2, mpeg1.1, mrc1b, mhc2, tlr3, tlr8b,* and *sting1*, whereas these signatures were markedly reduced in residual *irf8* mutant myeloid cells. Mutant cells instead showed increased expression of genes associated with immature myeloid states and immune responses, including *cd68, spi1b, pcna, s100a10b, lgals9l1, cd209l1, lect2.1, ifi44a1, irf8l1b,* and *irf9*, as well as genes involved in lysosomal function and intracellular trafficking, including *atp6ap2, ctsc, lamp2, lipf, npc2.2,* and *pikfyve*. Genes highlighted in red are analyzed in (**b-e**), Figures 4 and 6, and are shown here for reference within the broader transcriptional comparison. Dotted magenta boxes highlight genes with the largest changes based on expression level and fraction of cells expressing each gene.

**b** UMAP of reclustered macrophage/myeloid cells, showing segregation by genotype and diet. Representative mature LM genes (*c1ql1, cd74a, mhc2daa,* and *mpeg1.1*) were strongly enriched in WT cells but severely reduced in *irf8* mutant cells.

**c** UMAPs showing the distribution of genes enriched in residual *irf8* mutant myeloid cells, including *cd68, lgals9l1, npc2.2,* and *s100a10b*.

**d** Summary of transcriptional programs distinguishing WT LMs from residual *irf8* mutant myeloid cells from the liver scRNA-seq. WT cells retained a mature macrophage identity and functional program, whereas mutant cells showed reduced expression of these programs and increased expression of genes associated with immature myeloid states, immune responses, and lysosomal and intracellular trafficking functions.

**e** Representative single z-slices of 14 dpf WT and *irf8* mutant zebrafish livers showing distinct myeloid cell morphology and localization. WT LMs exhibit an elongated morphology with a few long branching processes and are closely aligned with the biliary network (yellow arrows). In contrast, residual *irf8* mutant myeloid cells are more amoeboid and preferentially localized around the vasculature (red arrows), and often contain endothelial-associated cargo. Few mutant cells can be associated with the biliary network, although most are not. These findings reveal a spatial redistribution of the small number of residual *irf8* mutant myeloid cells accompanying their transcriptional divergence from mature LMs.

**Supplementary Figure 10.**

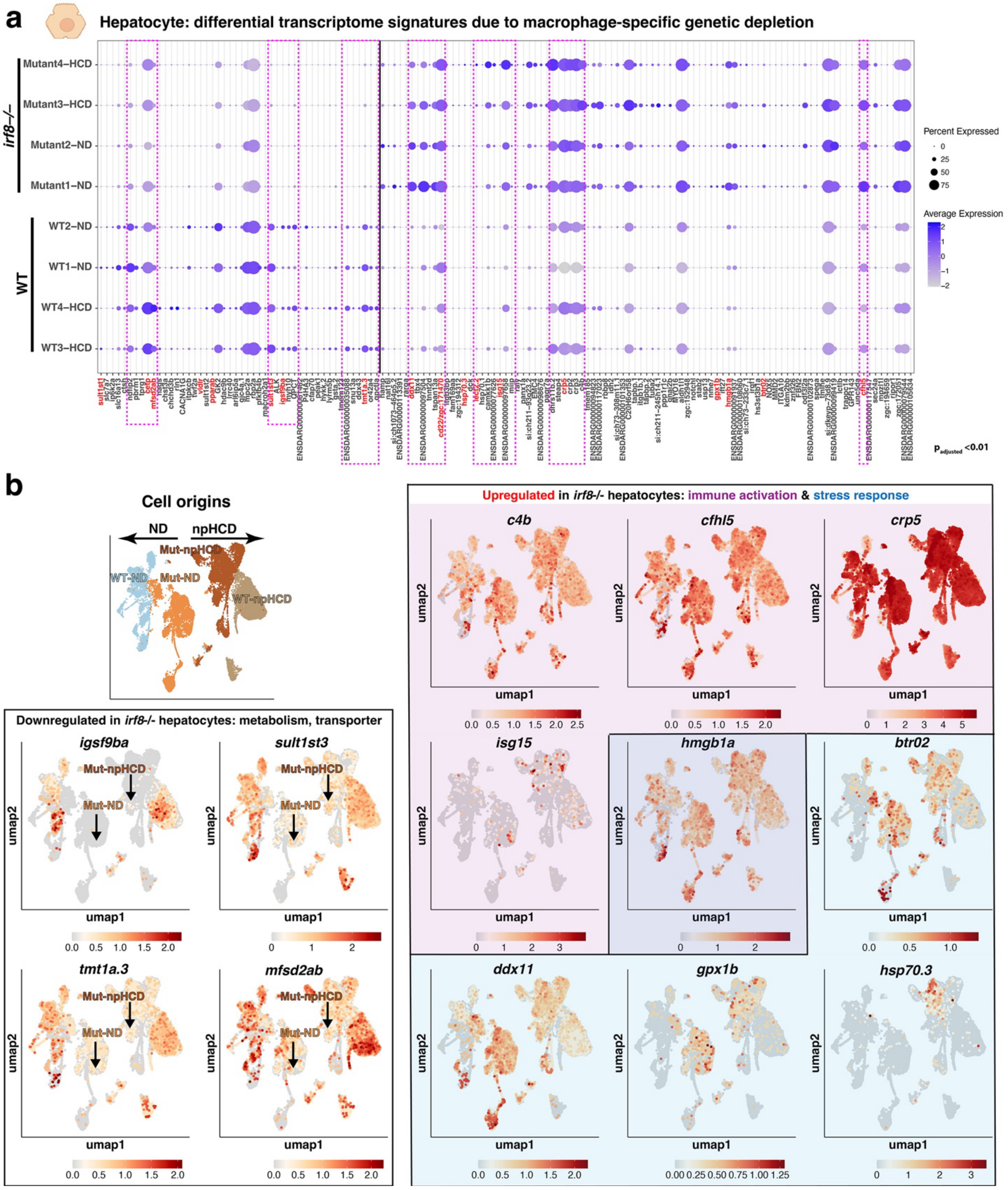

**Supplemental Figure 10 | Loss of liver macrophages reprograms hepatocytes toward adaptive immune- and stress- responsive states.**

**a** Bubble plot of differentially expressed genes (DEGs; adjusted $p < 0.01$) identified by unsupervised hierarchical clustering of hepatocytes reclustered from the liver scRNA-seq dataset (Figure 4). WT and *irf8* mutant hepatocytes segregated transcriptionally independent of diet (ND or npHCD). *irf8* mutant hepatocytes showed increased expression of immune-associated genes, including *c4b, cfhl5, crp* family members, *isg15, lect2.2, steap4, tapbp.1,* and *tapbp.2*; stress-associated genes, including *btr02, ddx11, gpx1b, hsp70.3, idh2, tmlhe,* and *usp13*; and cell-cycle-associated genes, including *ddx11, miip, raly,* and *pcna*. Conversely, mutant hepatocytes downregulated genes associated with metabolism and membrane lipid handling, including *sult1st3, mfsd2ab,* and *tmt1a.3*, as well as *igs9ba*. Genes highlighted in red were analyzed in **b** and Figures 4 and 6 and are shown here for reference within the broader transcriptional comparison. Dotted magenta boxes highlight genes with the largest changes based on expression level and fraction of cells expressing each gene.

**b** UMAPs of reclustered hepatocytes showing segregation by genotype and diet. Representative immune-associated genes (*c4b, cfhl5, crp5,* and *isg15*) and stress-associated genes (*btr02, ddx11, gpx1b,* and *hsp70.3*) were increased in *irf8* mutant hepatocytes, with *hmgb1a* spanning both transcriptional programs. Several immune- and stress-associated genes showed greater induction in mutant hepatocytes following npHCD. In contrast, *igs9ba, sult1st3, tmt1a.3,* and *mfsd2ab* showed reduced expression in mutant versus WT hepatocytes. The most significant gene expression changes between WT and *irf8* mutant hepatocytes occur across both diets, providing further support that the altered hepatocyte state is associated with LM loss itself, rather than a response to npHCD.

**Supplementary Figure 11.**

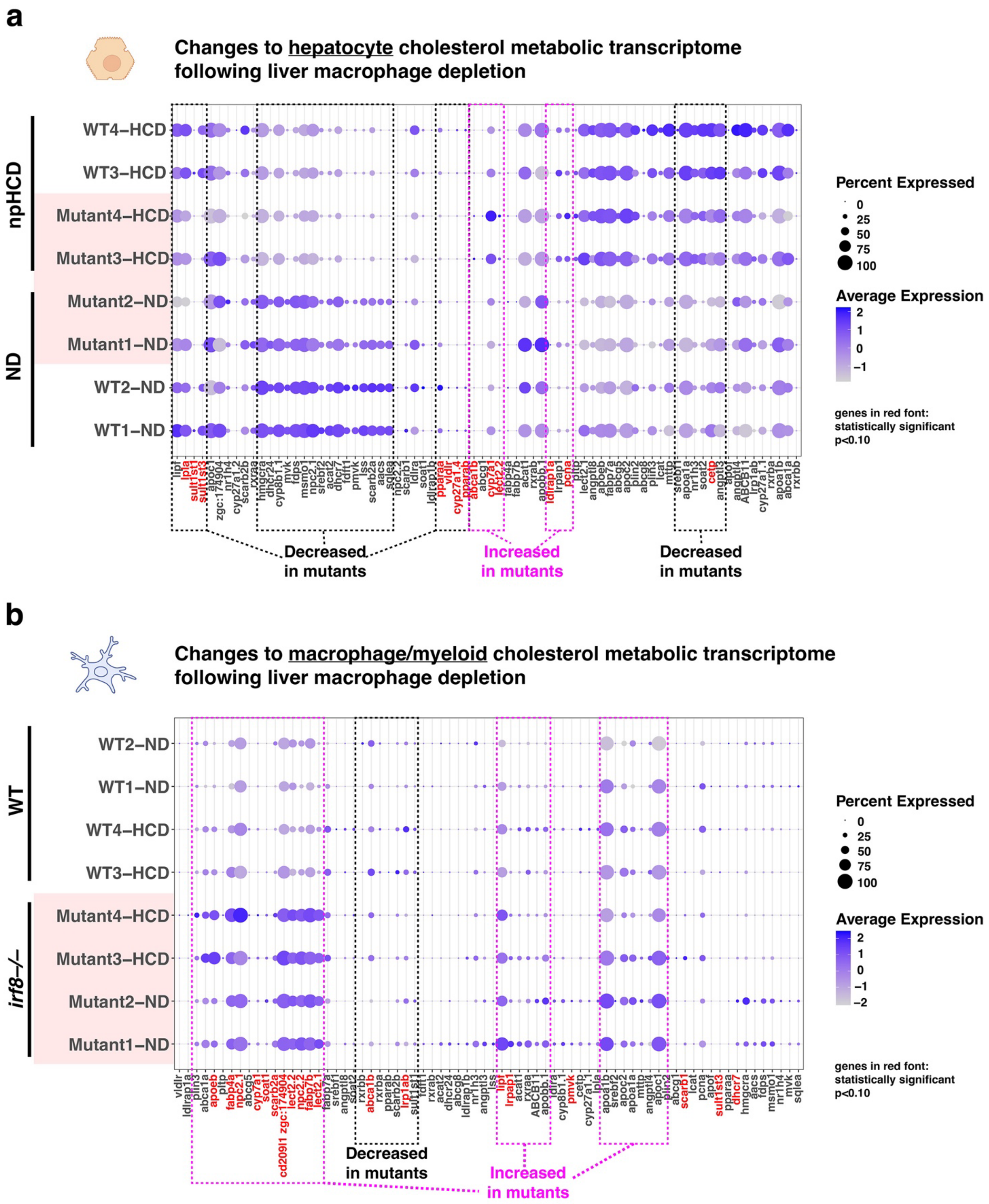

**Supplemental Figure 11 | Loss of mature liver macrophages reshapes cholesterol-handling programs in hepatocytes and residual myeloid cells.**

**a** Bubble plot with unsupervised hierarchical clustering of reclustered hepatocytes from the liver scRNA-seq dataset (Figure 4), showing expression of 75 genes with established roles in cholesterol metabolism across WT and *irf8* mutant larvae fed ND or npHCD. Hierarchical clustering primarily separated hepatocytes by dietary condition. Relative to WT, *irf8* mutant hepatocytes increased expression of *cyp7a1*, the rate-limiting enzyme in classical bile acid synthesis, and *abca1b*, a mediator of cellular cholesterol efflux, while reducing expression of cholesterol biosynthesis genes (*hmgcra, sqlea*), the lipoprotein-remodeling gene *cetp*, sulfotransferases (*sult1st3* and *sult1st1*), and the bile acid receptor *nr1h4* (FXR). This coordinated shift from cholesterol synthesis toward bile acid production and efflux is consistent with an adaptive response to increased hepatocyte cholesterol burden following loss of mature LMs.

**b** Bubble plot with unsupervised hierarchical clustering of reclustered macrophages/myeloid cells from the liver scRNA-seq dataset (Figure 4), showing 73 detectable cholesterol metabolism-associated genes across WT LMs and residual *irf8* mutant myeloid cells under ND and npHCD. In contrast to hepatocytes, these populations clustered primarily by genotype, revealing distinct cholesterol-handling programs in mature WT LMs versus residual *irf8* mutant myeloid cells. Residual mutant cells showed increased expression of genes associated with intracellular cholesterol trafficking and transport together with reduced expression of cholesterol export *abca1b*, consistent with altered intracellular cholesterol handling and impaired cholesterol efflux. Genes highlighted in red met the significance threshold ($p < 0.10$). Dotted boxes highlight gene clusters showing prominent genotype-associated changes in at least one dietary condition.

**Supplementary Figure 12.**

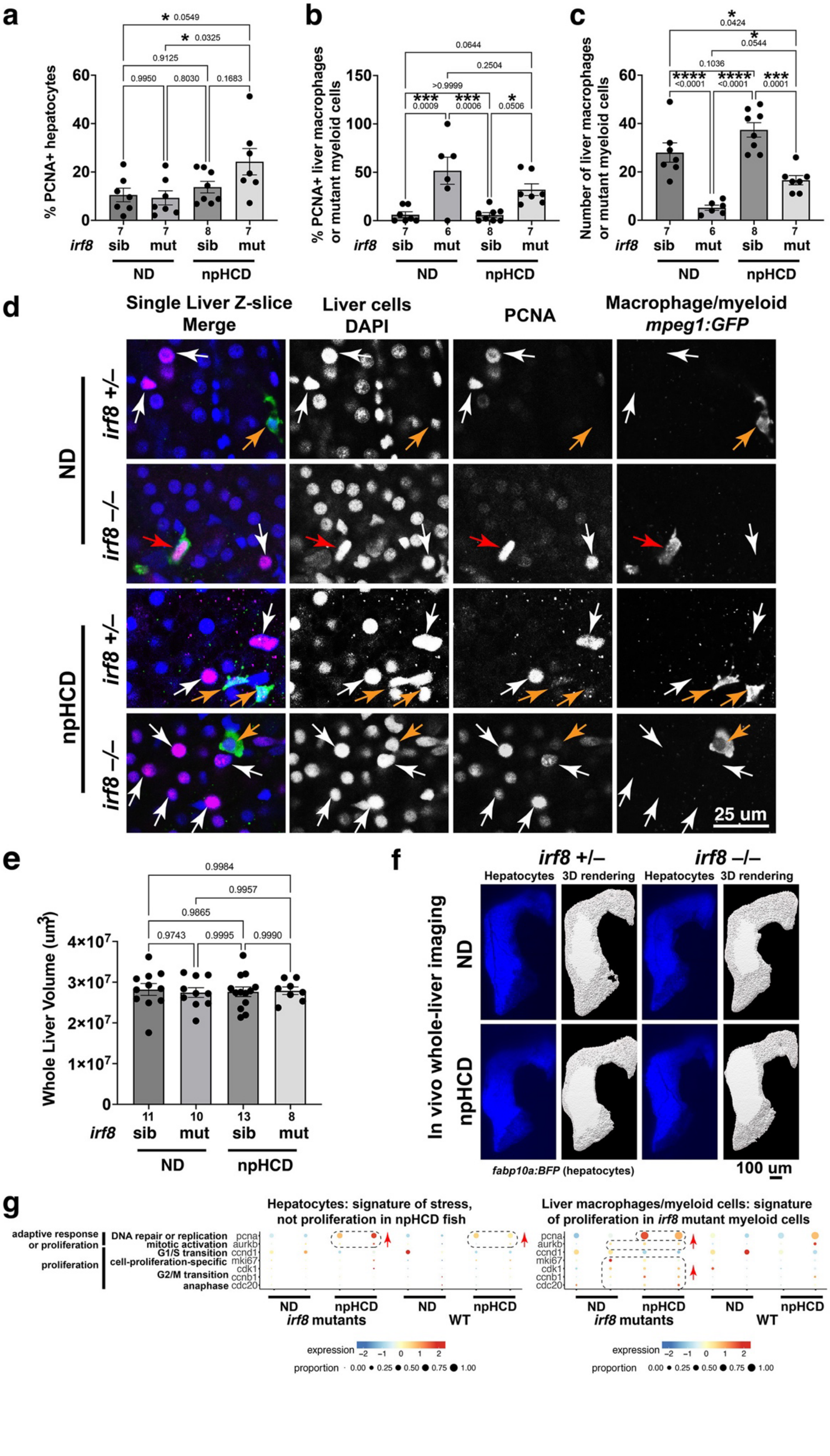

**Supplemental Figure 12 | Loss of mature liver macrophages induces an adaptive stress-associated replication response in hepatocytes without overt proliferation or changes in liver size.**

**a** Quantification of $PCNA^+$ hepatocytes in WT and *irf8* mutant larvae following ND or npHCD feeding. *irf8* mutants showed a modest, non-significant increase in $PCNA^+$ hepatocytes after npHCD compared with WT, whereas npHCD significantly increased the proportion of $PCNA^+$ hepatocytes in *irf8* mutants relative to ND.
**b** Quantification of $PCNA^+$ liver macrophages or residual myeloid cells. *irf8* mutant livers exhibited a significant increase in $PCNA^+$ residual myeloid cells irrespective of diet, indicating enhanced DNA replication associated with loss of *irf8*-dependent mature LM identity.
**c** Quantification of total liver macrophage/myeloid cell number. The markedly reduced myeloid cell population persisted in *irf8* mutant livers across conditions, but npHCD significantly increased the number of residual mutant myeloid cells. Together with the increased $PCNA^+$ fraction, these data support an npHCD-associated increase in residual myeloid-cell proliferation.
**d** Representative confocal images of PCNA immunostaining in WT and *irf8* mutant livers following ND or npHCD feeding. *mpeg1* cells were detected with anti-GFP, nuclei with DAPI, and DNA replication with anti-PCNA. White arrows indicate $PCNA^+$ hepatocytes; orange arrows indicate $DAPI^+$ nuclei within *mpeg1*$^+$ cells with weak or undetectable PCNA; red arrows indicate $PCNA^+$ *mpeg1*$^+$ macrophage/myeloid cells.
**e** Quantification of whole-liver volume in WT and *irf8* mutant larvae following ND or npHCD feeding. Neither genotype nor diet altered liver size.
**f** Representative confocal volumetric images of whole livers in live zebrafish imaged with a 20× objective (left) and corresponding 3D Imaris surface reconstructions (right). *fabp10a:BFP* labels the liver for volumetric analysis.
**g** Bubble plots showing selected genes associated with DNA replication/cellular stress, and proliferation in hepatocytes and liver macrophage/myeloid cells. npHCD increased *pcna* and *aurkb* expression in hepatocytes of both genotypes without induction of the broader canonical proliferation program. In contrast, residual *irf8* mutant myeloid cells showed increased *pcna* together with canonical cell-cycle/proliferation genes relative to WT LMs, consistent with a more proliferative state.
Data are mean ± SEM. Each point represents one larva; *N* shown below bar graphs. Statistical significance was assessed by one-way ANOVA followed by multiple pairwise comparisons. All data generated in this figure came from 14 dpf zebrafish after 5-day diet feeding.

**Supplementary Figure 13.**

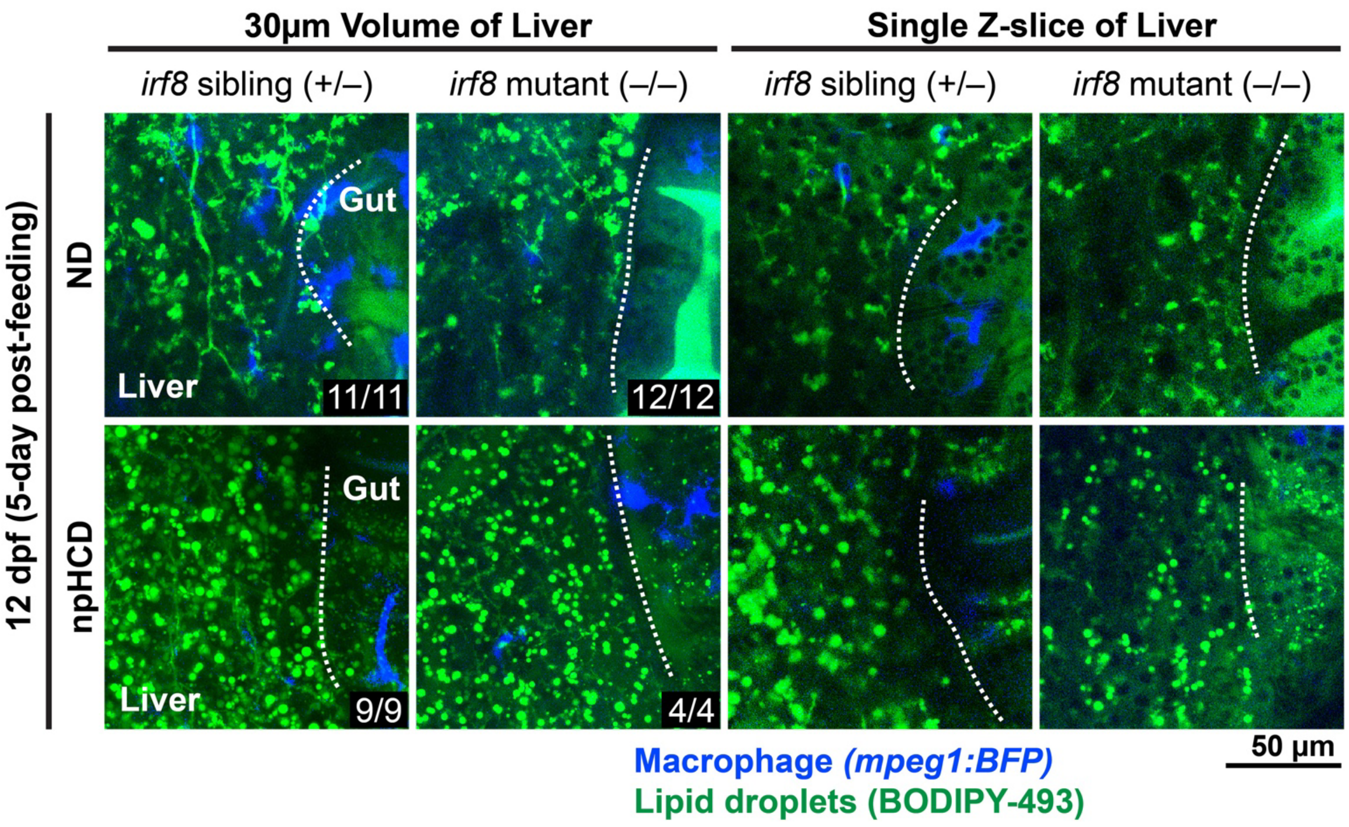


**Supplemental Figure 13 | High-cholesterol feeding drives neutral lipid accumulation independently of macrophage status.**

Representative confocal images of BODIPY-493 staining in 12 dpf *irf8* sibling and *irf8* mutant larvae following 5 days of normal diet (ND) or non-pathological high-cholesterol diet (npHCD) starting at 7 dpf. Images show 30-µm maximum-intensity projections and representative single z-slices of the liver and proximal gut. BODIPY-493 staining revealed increased neutral lipid droplet accumulation following npHCD feeding in both genotypes, with no apparent genotype-dependent differences in the liver or proximal gut. ND-fed larvae exhibited markedly fewer lipid droplets in both tissues. Numbers in the lower right indicate the number of larvae analyzed that exhibited the same pattern as the representative image.

**Supplementary Figure 14.**

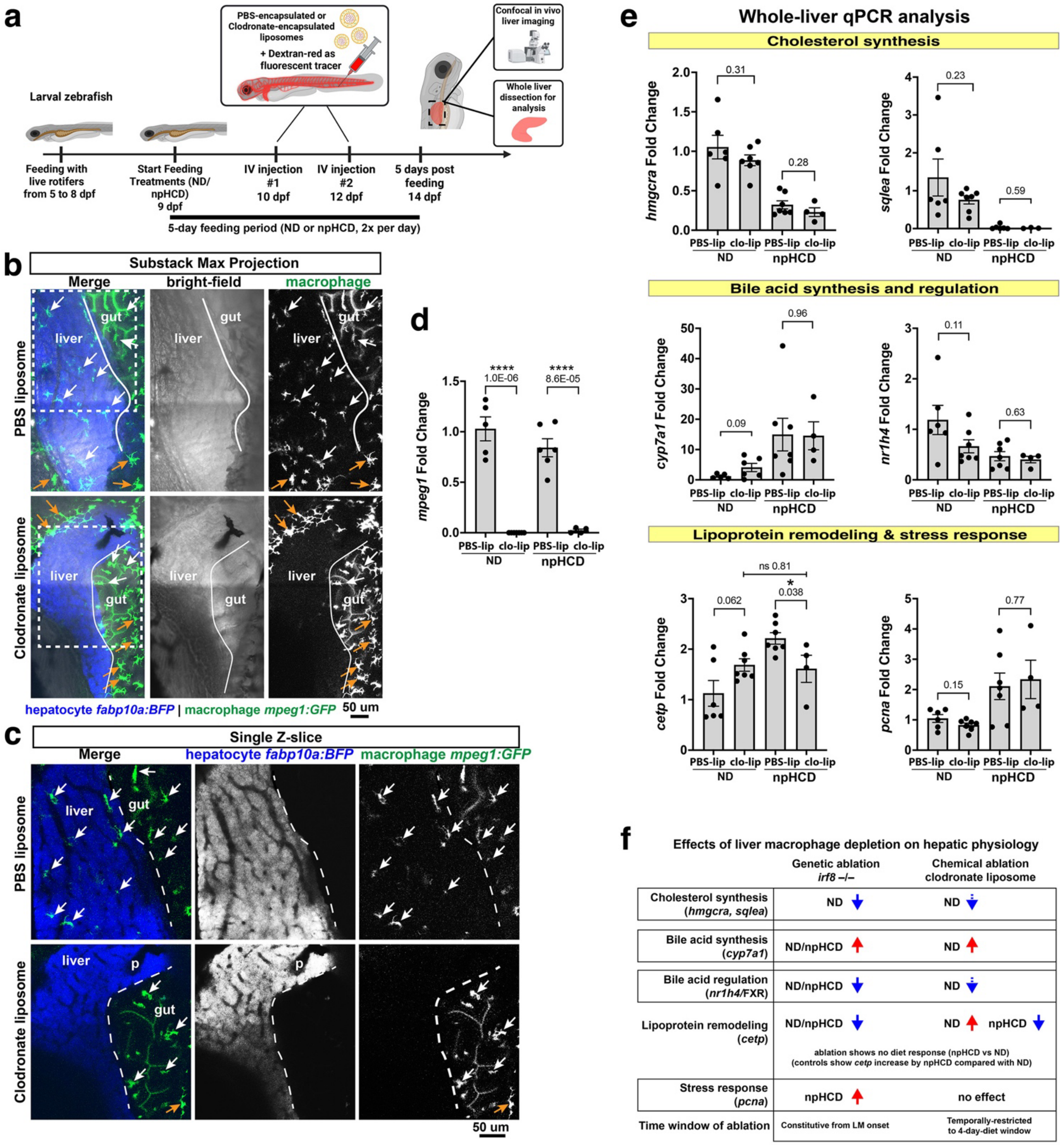

**Supplemental Figure 14 | Acute clodronate-mediated depletion of liver macrophages alters hepatic cholesterol-associated transcriptional responses.**

**a** Schematic of the experimental strategy for acute liver macrophage depletion using clodronate-containing liposomes. Larvae maintained on ND or npHCD received intravenous injections of PBS-containing control liposomes or clodronate-containing liposomes at 10 and 12 dpf, allowing liver macrophages to establish before depletion.
**b** Representative confocal images of the digestive tract, including liver and gut, in *fabp10a:mpeg1* larvae following PBS- or clodronate-liposome treatment. Clodronate treatment markedly targeted and depleted *mpeg1*$^{+}$ macrophages from the liver while largely preserving the intestinal macrophage population. PBS-treated controls retained abundant liver and intestinal macrophages.
**c** Higher-magnification views of the regions shown in **b**, highlighting the selective loss of liver macrophages following clodronate treatment while intestinal macrophages appear intact. Arrows indicate liver- or gut-resident macrophages; dashed lines delineate the liver–gut boundary.
**d** qPCR analysis of the macrophage marker *mpeg1* showing significantly reduced expression following clodronate-mediated depletion, confirming effective reduction of the liver macrophage population.
**e** qPCR analysis of six cholesterol metabolism-associated genes selected based on transcriptional changes in macrophage-depleted *irf8* mutants. Acute clodronate-mediated depletion produced transcriptional trends that broadly paralleled those observed following genetic loss of mature liver macrophages, although individual changes did not reach statistical significance.
**f** Summary of hepatic cholesterol-associated transcriptional responses following chronic genetic loss versus acute depletion of mature liver macrophages. Notably, npHCD normally induces *cetp* expression, whereas this dietary response was blunted following either genetic or clodronate-mediated macrophage depletion in part due to a significant reduction in hepatic cetp expression after macrophage loss in both depletion paradigms. The concordant loss of *cetp* induction across two independent depletion paradigms supports a role for liver macrophages in coordinating the hepatic transcriptional response to dietary cholesterol.
Data are mean ± SEM. Statistical significance was determined as indicated in the figure.

**Supplementary Figure 15.**

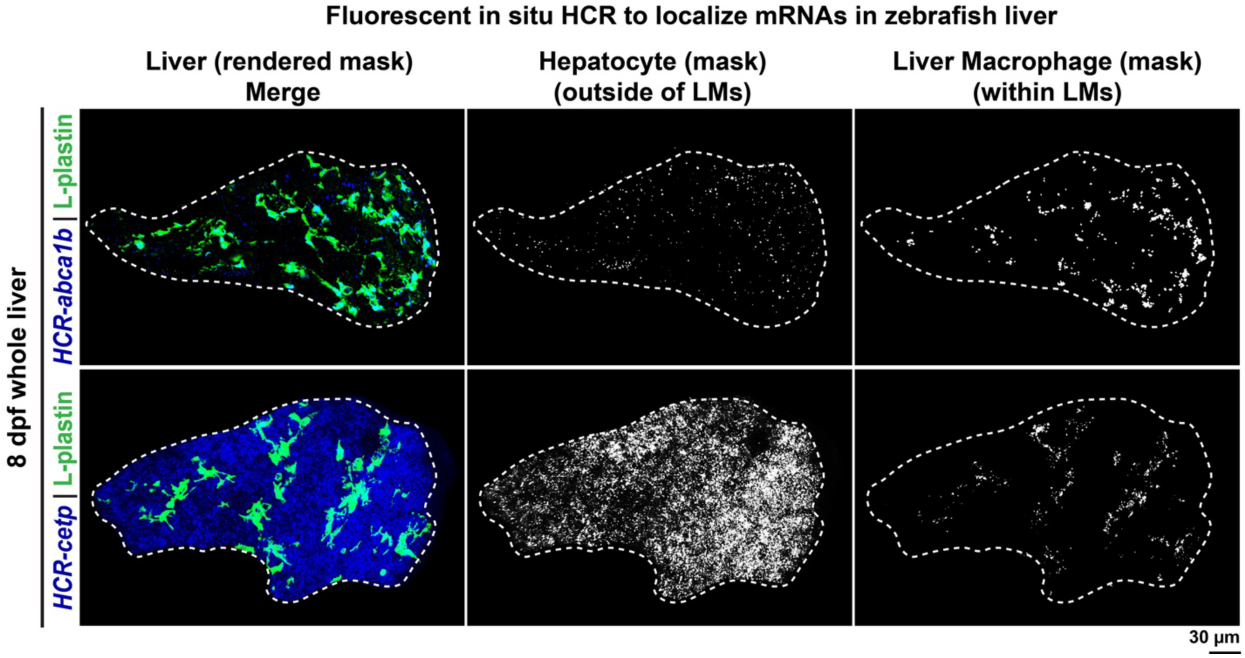


**Supplementary Figure 15 | RNA in situ HCR reveals macrophage restricted *abca1b* expression but broad *cetp* expression in the zebrafish larval liver.**

Fluorescent in situ hybridization chain reaction (HCR) was used to localize *abca1b* and *cetp* mRNAs in the larval zebrafish liver. Anti-L-plastin immunostaining was performed concurrently to identify hepatic macrophages and distinguish them from other liver cells. Confocal volumetric images show merged HCR (blue) and L-plastin (green) signals within an Imaris-generated liver mask, which excludes fluorescence outside the liver. Right panels show separate masks of L-plastin$^{+}$ liver macrophages (LMs) and L-plastin$^{-}$ hepatic cells, enabling direct assessment of cell-type-specific RNA expression. *abca1b* expression was highly enriched in L-plastin$^{+}$ LMs, whereas *cetp* expression was broadly distributed among LMs and other hepatic cells, supporting distinct cellular sources for these cholesterol-handling genes.

**Supplementary Figure 16.**

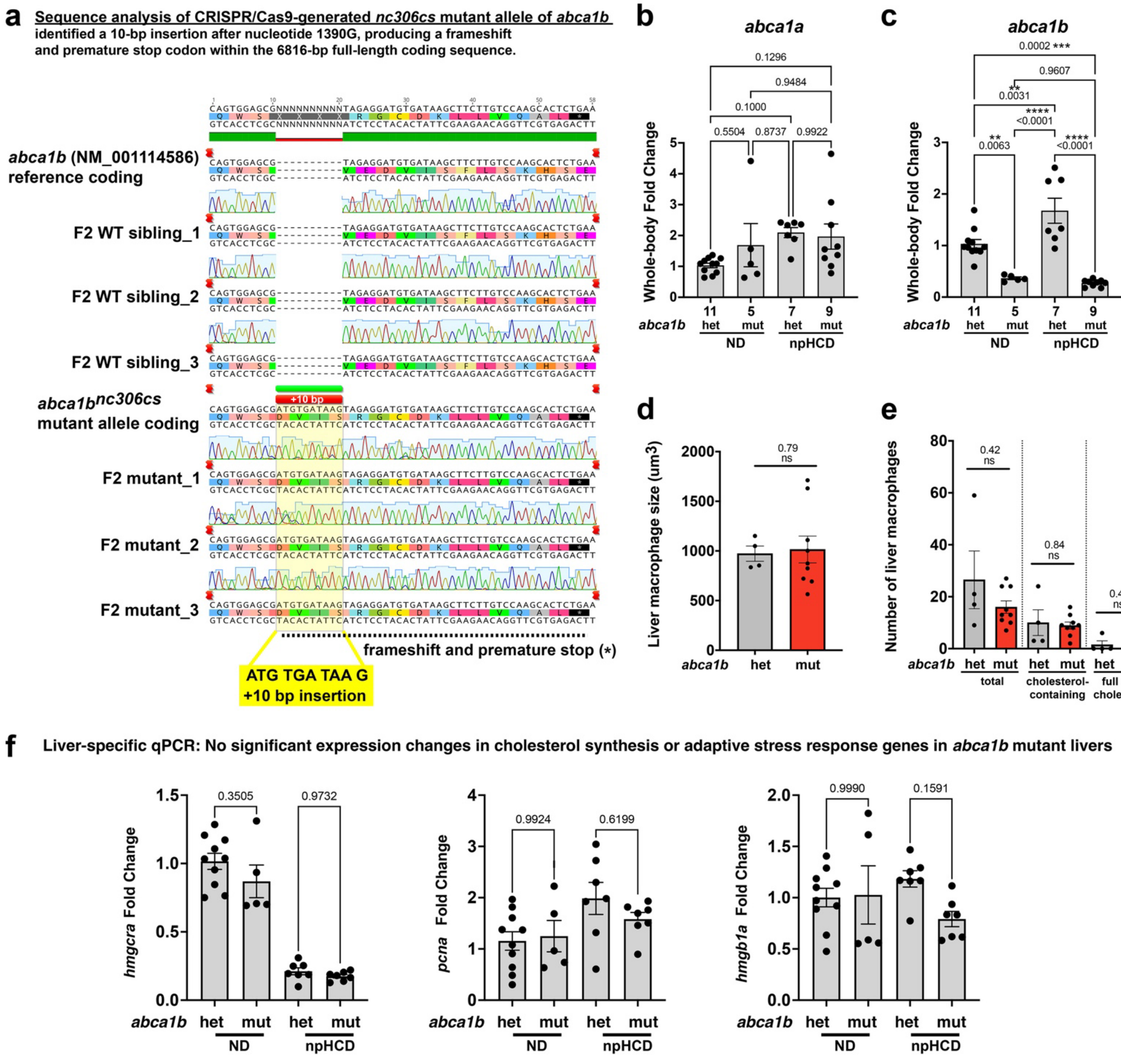

**Supplemental Figure 16. Identification and characterization of CRISPR-generated *abca1b*(nc306cs) mutant zebrafish.**

**a** Sanger sequencing identifies a 10-bp insertion in *abca1b*(nc306cs), resulting in a frameshift and premature stop codon within the 6,816-bp coding sequence, predicted to generate a loss-of-function allele. Alignment shows DNA sequence at the targeted locus from 3 independent WT siblings and 3 independent mutant samples showing the 10-bp insertion specifically in the mutants.
**b,c** Relative mRNA expression of *abca1a* (**b**) and *abca1b* (**c**) measured by qPCR in whole-body samples from *abca1b* heterozygous and mutant larvae. *abca1a* expression was unchanged, indicating no detectable transcriptional compensation, whereas *abca1b* expression was significantly reduced in mutants, consistent with the predicted loss-of-function allele.
**d** Quantification of liver macrophage volume showed no difference between *abca1b* heterozygous and mutant macrophages, indicating that *abca1b* loss does not alter LM size.
**e** Although the fraction of LMs exhibiting circulating cholesterol uptake was significantly altered in *abca1b* mutants (Figure 6), the total number of LMs and the number of LMs within each cholesterol-uptake category were unchanged. Thus, the altered uptake phenotype reflects a shift in the distribution of cholesterol uptake among LMs rather than a change in LM abundance.
**f** qPCR analysis of cholesterol- and stress-associated genes altered following LM depletion in *irf8* mutants. Expression of *hmgcra, pcna,* and *hmgb1a* was unchanged in *abca1b* mutants, indicating that *abca1b* loss does not fully recapitulate the broader cholesterol-associated transcriptional and stress response induced by LM depletion.
Data are mean ± SEM. Each point represents one animal (**b,c**) or one liver (**d–f**); numbers of biological replicates are indicated below the bar graphs. Statistical significance was determined by one-way ANOVA followed by multiple comparisons (**b,c**) or two-tailed Student's *t*-test (**d–f**).

**Supplementary Figure 17.**

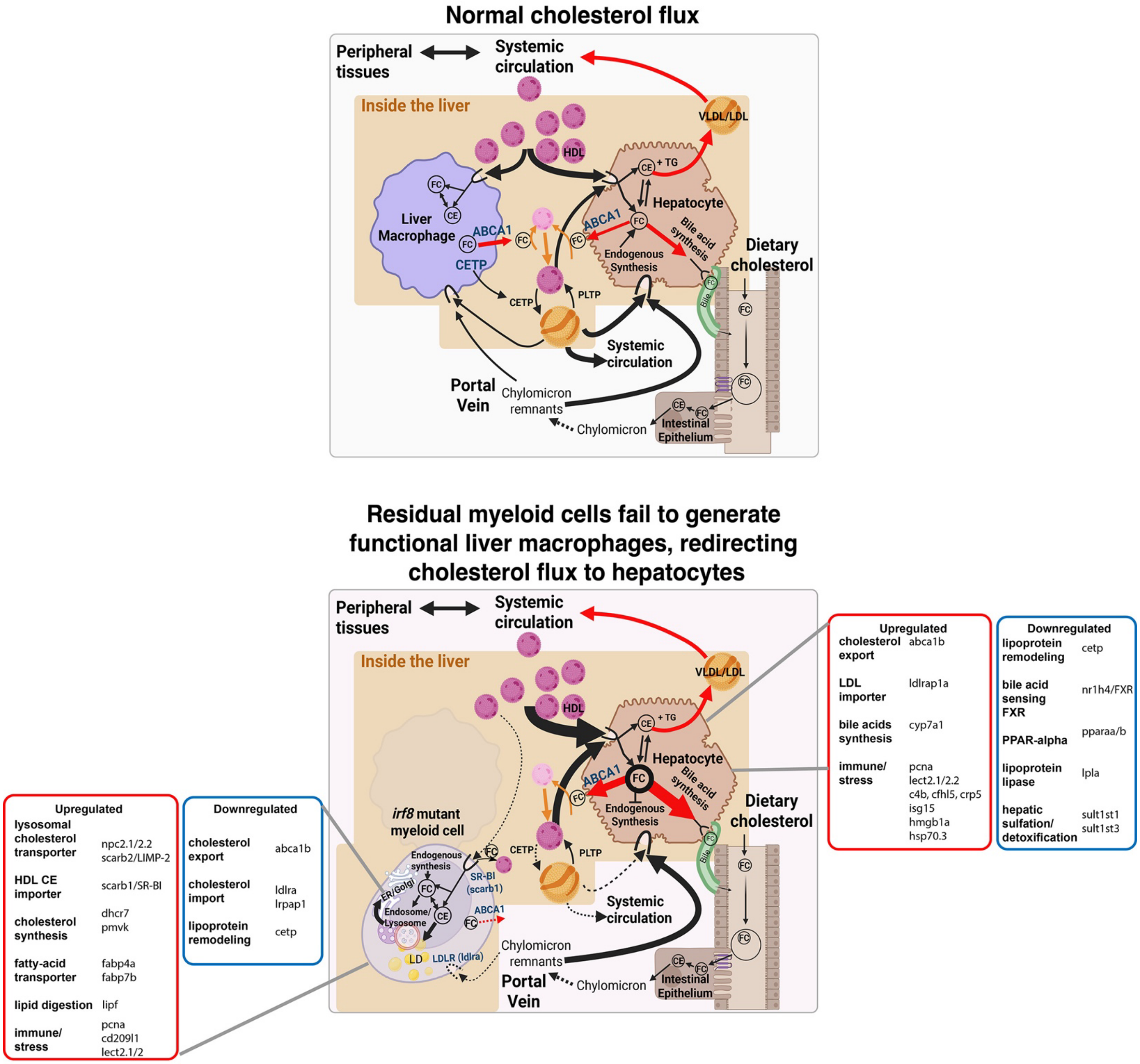

**Supplemental Figure 17 | Working model: Liver macrophages emerge as cellular buffers regulating cholesterol flux to ensure hepatic homeostasis.**

Schematic illustrating the proposed role of mature liver macrophages (LMs) in hepatic cholesterol homeostasis and the consequences of their loss. Under homeostatic conditions, mature LMs form a specialized hepatic compartment positioned at the interface of the biliary and vascular networks. This strategic localization enables LMs to access circulating and hepatocyte-derived cholesterol, internalize and transiently store cholesterol, promote cholesterol efflux through ABCA1, and contribute to lipoprotein remodeling through CETP. By partitioning and buffering hepatic cholesterol flux, LMs may limit fluctuations in hepatocyte cholesterol burden and thereby support coordinated cholesterol and bile acid homeostasis.

Following loss of mature LMs, residual *irf8* mutant myeloid cells fail to acquire the transcriptional identity, spatial organization, and cholesterol-handling programs characteristic of mature LMs. We propose that loss of this macrophage buffering compartment redistributes cholesterol flux toward hepatocytes, which respond by shifting cholesterol handling toward efflux and disposal, including increased ABCA1-associated export and bile acid synthesis, together with reduced cholesterol biosynthesis and lipoprotein-remodeling programs. This altered cholesterol state is accompanied by hepatocyte stress- and replication-associated transcriptional responses. Significant transcriptional changes identified in residual myeloid cells and hepatocytes in LM-depleted *irf8* mutants are highlighted. Together, the model positions mature LMs as an active cellular buffer and regulator of hepatic cholesterol flux, linking circulating cholesterol handling to hepatocyte metabolic and stress responses.

**Supplementary Movie 1 | Large-volume, low-density rearing in 1-L tanks with live and dry diets promotes active swimming and feeding for normal maturation of larval zebrafish.**

Larvae were maintained in a stand-alone incubator under a 14-h light/10-h dark cycle at 28.5°C, conditions optimized for zebrafish growth in 1-L tanks. The 1-L tank volume and vertical swimming space allow larvae to actively swim throughout the tank, rather than being confined to the shallow dimensions of a Petri dish, while the low fish density (> 20 mL per larva) and high water-to-fish ratio provide an environment that supports larval maturation. Following introduction of dry food, 12 dpf larvae actively swim toward and congregate at the water surface to feed. The video covers 8-second of zebrafish activity and is shown in real time immediately after dry food was sprinkled onto the water surface. See also Supplementary Fig. 1.

**Supplementary Movie 2 | Fly-through of a confocal z-stack showing the spatial localization of liver-resident macrophages (LMs) relative to the biliary and vascular networks in a WT 12 dpf zebrafish liver.**

In vivo confocal imaging was used to localize macrophages (*mpeg1:BFP*) relative to the biliary network (*krt18:GFP*) and vasculature (*kdrl:mCherry*) throughout larval development, shown here in a 12 dpf larva. The movie traverses a 78 μm sub-z-stack and reveals LMs (few examples, magenta arrows) intimately intertwined with the biliary epithelial cells and intimately overlapping the biliary network throughout the liver volume. In contrast, LM cell bodies are neither within nor closely associated with the endothelial vasculature, although their cellular processes can extend to and contact blood vessels. Images were acquired at 3 μm z-intervals using a 40× objective from a 1 × 2 tile scan at 1024 × 1024 pixels per tile. Imaging extended from the skin (white arrow) overlying the liver to the deepest detectable region of the liver. As expected from the reporters used, peripheral macrophages ($BFP^+$) and some superficial skin cells ($GFP^+$) are also visible outside the liver. The movie shows the zebrafish with anterior to the right and ventral side at the bottom, rotated 90° relative to the orientation shown in Fig. 1e.

**Supplementary Movie 3 | Liver macrophages intimately contact and extend around and into the hepatic vasculature.**

3D visualization of an approximately 15 μm confocal z-stack acquired in vivo from the liver of a 14 dpf WT zebrafish expressing *mpeg1:GFP* to label macrophages, *fabp10a:BFP* to label hepatocytes, and *kdrl:mCherry* to label the vasculature. The movie shows three 360° rotations, two around the y-axis and one around the x-axis, to reveal the intricate spatial relationships between LMs and blood vessels. Fine LM processes extend into and around the vasculature at multiple sites, while a centrally positioned LM has a substantial branch of its cell body that crosses into the sinusoidal lumen. Images were acquired using a 40× objective. See also the corresponding Supplementary Fig. 3a.

**Supplementary Movie 4 | Liver macrophages dynamically sample inside the hepatic blood vessels.**

In vivo time-lapse confocal imaging show dynamic hair-like LM processes extend and retract within the vessel, while major cellular branches can penetrate the sinusoidal lumen in the 8 dpf zebrafish liver. Three transgenes were used: *mpeg1:GFP* for macrophages, *kdrl:mCherry* for vasculature, and *fabp10a:BFP* for hepatocytes. Confocal z-stack was captured at 3 μm z-intervals continuously at every ~24 seconds using a 40× objective at 1024 × 1024 pixels per image. Total imaging time was 30 minutes and 50 seconds. Time stamp shows h:mm:ss. See also the corresponding Supplementary Fig. 3b.

**Supplementary Movie 5 | Liver macrophages are anchored along the biliary network.**

In vivo time-lapse confocal imaging shows that liver-resident macrophages (LMs, white arrows) maintain stable positions along the biliary network, in striking contrast to the dynamic migration of peripheral macrophages (dotted white arrows). LMs remain closely associated and aligned with the biliary epithelial cells, revealing a distinct stationary behavior characteristic of their resident niche. Three transgenes were used: *mpeg1:GFP* for macrophages, *kdrl:mCherry* for vasculature, and *krt18:GFP* for biliary epithelial cells (which also labels a subset of skin cells in the periphery). Images came from time-lapse confocal z-stacks captured at 3 μm z-intervals every 10 minutes using a 40× objective from a 1 × 2 tile scan at 512 × 512 pixels per tile in a live 10 dpf zebrafish. Total imaging time was 3 hours. Time stamp shows h:mm:ss. See also the corresponding Supplementary Fig. 4.

**Supplementary Movie 6 | Liver macrophages dynamically capture circulating tracers.**

Time-lapse spinning-disk confocal imaging of a live 7 dpf zebrafish expressing the macrophage reporter *mpeg1:BFP* after intravenous injection of Alexa488-conjugated LPS. Circulating fluorescent LPS reveals hepatic blood flow and the vascular network in real time. LMs dynamically interact with the circulation, with some accumulating substantial tracer (asterisk) and others extending processes to sense and capture smaller amounts of circulating LPS (arrows). High-magnification imaging reveals spine-like protrusions enclosing circulating particles and dynamic processes extending around and into blood vessels. Images were acquired at 3 μm z-intervals every ~15 s for ~2 h using a 40× objective, shown at 10 fps. Timestamp, h:mm:ss. See also Fig. 3a for selected stills from a cropped time series of a single LM (red arrow).
